# PP2A Dephosphorylates Phytochrome-Interacting Factor 3 to Modulate Photomorphogenesis in *Arabidopsis*

**DOI:** 10.1101/2023.08.17.553727

**Authors:** Xingbo Cai, Sanghwa Lee, Andrea Gomej-Jaime, Wenqiang Tang, Yu Sun, Enamul Huq

## Abstract

The phytochrome (phy) family of sensory photoreceptors modulates developmental programs in response to ambient light. phys control gene expression in part by directly interacting with the bHLH class of transcription factors, phytochrome-interacting factors (PIFs), and inducing their rapid phosphorylation and degradation. Several kinases have been shown to phosphorylate PIFs and promote their degradation. However, the phosphatases that dephosphorylate PIFs are less investigated. Here we describe the identification of four regulatory subunits of *Arabidopsis* protein phosphatase 2A (PP2A) family (B’α, B’β, B’’α and B’’β) that interacted with PIF3 in yeast-two-hybrid, *in vitro* and *in vivo* assays. The *pp2ab’’αβ* and *b’’αβ/b’αβ* displayed short hypocotyls, while the overexpression of the B subunits induced longer hypocotyls compared to wild type under red light. The light-induced degradation of PIF3 was faster in *b’’αβ/b’αβ* quadruple mutant compared to wild type. Consistently, immunoprecipitated PP2A A and B subunits directly dephosphorylated PIF3-MYC *in vitro*. RNA-seq analyses showed that B’’α and B’’β alter global gene expression in response to red light. *PIFs (PIF1, PIF3, PIF4* and *PIF5)* are epistatic to these four B subunits in regulating hypocotyl elongation under red light. Collectively, these data show an essential function of PP2A in dephosphorylating PIF3 to modulate photomorphogenesis in *Arabidopsis*.

## Introduction

Environmental factors such as light has profound impacts on all organisms. This is especially important for plants due to their sessile nature. Plants employ a battery of sensory photoreceptors for perceiving and responding to surrounding light environment (Bae and Choi, 2008). The phytochrome (phy) family encoded by five members in Arabidopsis (PHYA-PHYE) perceives and responds to the red/far-red region of the light spectrum and controls development throughout plant life cycle (Legris et al., 2019; Cheng et al., 2021). Upon red light exposure, phys change conformation from the inactive Pr form to the light activated Pfr form. The activated Pfr form is translocated from the cytoplasm into the nucleus and interacts with multiple nuclear proteins. Among those, the Phytochrome-Interacting Factors (PIFs), a small family of basic helix-loop-helix (bHLH) transcription factors, are primary interacting partners as they specifically bind to the Pfr form (Leivar and Quail, 2011; Pham et al., 2018a). PIFs function primarily as negative regulators of photomorphogenesis to repress seed germination, promote hypocotyl elongation and shade avoidance responses (Lee and Choi, 2017; Pham et al., 2018a). To remove these negative regulators, light-activated phys induce rapid phosphorylation, ubiquitination and degradation of PIFs through the 26S proteasome pathway to promote photomorphogenesis (Pham et al., 2018a; Cheng et al., 2021). In addition, phys also interact with PIFs and inhibit the DNA binding and transcriptional activation activity of PIFs in response to light (Park et al., 2012; Park et al., 2018; Yoo et al., 2021).

In recent years, multiple kinases and E3 ubiquitin ligases involved in PIF degradation have been described (Cheng et al., 2021). SUPPRESSOR OF PHYA-105 1 family members (SPA1-SPA4) directly interact with E3 ligase CONSTITUTIVELY PHOTOMORPHOGENIC1 (COP1) and function as a cognate kinase-E3 ligase complex to mediate light-induced degradation of PIF1 to promote seed germination and seedling development (Zhu et al., 2015; Pham et al., 2018b; Paik et al., 2019). Casein Kinase 2 (CK2) directly phosphorylates PIF1 in a light-independent manner. However, CK2-mediated phosphorylation of PIF1 is necessary for the light-induced degradation of PIF1 (Bu et al., 2011). Photoregulatory Protein Kinases (PPK1-4) interact with and phosphorylates PIF3 in a light-induced fashion (Ni et al., 2017) and induce PIF3-phyB co-degradation. In addition, MPK6 has been shown to phosphorylate PIF3 and regulates its turnover (Xin et al., 2017). Interestingly, SALT OVERLY SENSITIVE2 (SOS2) phosphorylates PIF1 and PIF3 to promote their degradation under salt stress conditions, while it phosphorylates PIF4 and PIF5 to stabilize them under shade conditions (Han et al., 2023; Ma et al., 2023). However, PIFs are not only degraded in the light but also under dark conditions. The GSK3-like kinase BRASSINOSTEROID-INSENSITIVE (BIN2) phosphorylates both PIF3 and PIF4 and promotes their degradation in dark conditions (Ling et al., 2017).

Reversible phosphorylation of phytochromes as well as phy signaling partners plays crucial roles in light signaling pathways. For example, phosphorylation of phytochromes regulates their interaction with signaling partners (Choi et al., 1999; Kim et al., 2004; Nito et al., 2013), as well as dark reversion of phyB (Medzihradszky et al., 2013). The catalytic subunit of an *Arabidopsis* protein phosphatase PP2A (AtFyPP3) interacts with, and dephosphorylates oat phyA to regulate flowering time (Kim et al., 2002). A type 5 protein phosphatase (PAPP5) specifically dephosphorylates the Pfr form of phytochromes and enhances phytochrome signaling (Ryu et al., 2005). Phytochrome-associated protein phosphatase type 2C directly dephosphorylates phytochromes and indirectly mediates the *in vitro* dephosphorylation of PIF3 (Phee et al., 2008). Only two phosphatases, TOPP4 (catalytic subunit of PP1) and Fypp1/Fypp2 (catalytic subunits of PP6), have been shown to dephosphorylate PIF3-PIF5 and regulate their abundance (Yue et al., 2016; Yu et al., 2019). However, Fypp1/2 mainly functions in the dark to repress photomorphogenesis. Although, TOPP4 dephosphorylates the light-induced phosphorylated form of PIF5, TOPP4 mutant has been isolated as a dwarf mutant, suggesting a more general function in regulating seedling growth. Thus, the phosphatase(s) that is/are functioning as a *bona fide* light signaling component is still unknown.

PP2A is a group of ubiquitous and highly conserved serine-threonine phosphatases, involved in the growth and development and phytohormone signaling pathways in plant (Luan, 2003; Farkas et al., 2007; Booker and DeLong, 2016; Bheri et al., 2021). In *Arabidopsis*, functional PP2A is a heterotrimeric complex, consisting of a scaffolding A subunit (3 isoforms) and a catalytic C subunit (5 isoforms), and a regulatory B subunit (18 isoforms). The A and C subunits act as core enzyme, while the B subunit determines substrate specificity, subcellular localization of the ABC trimer. The catalytic subunits of PP2A and other PPP family members are conserved due to the high sequence similarity. Also, the three A subunits show similar functions, because of their high amino acid sequence similarity. Unlike the conserved A and C subunits, the members of the B subunit exist in different gene families. In *Arabidopsis*, B subunits are further divided into B, B’, and B’’ subfamilies. The spatiotemporal gene expression and subcellular localization of these subunits are involved in determining the substrate specificity and enzymatic activity of PP2A family (Farkas et al., 2007).

In an effort to isolate phosphatase (s) that is/are involved in dephosphorylation of the light-induced form of PIFs, we identified four regulatory subunits of PP2A (B’α, B’β, B’’α and B’’β) that interact with PIF3 *in vitro* and *in vivo*. We show that B’α, B’β, B’’α and B’’β delay the red-light induced PIF3 degradation and promote hypocotyl elongation specifically under red-light condition. Thus, PP2A regulatory subunits act as new negative regulators of phyB signaling by dephosphorylating PIF3 and preventing PIF3 degradation.

## Results

### PIF3 interacts with PP2A B’α, B’β, B’’α and B’’β *in vivo* and *in vitro*

To identify the potential phosphatases that interact with PIFs, we performed a yeast-2-hybrid (Y2H) screening by using PIF3 as a bait. We identified four regulatory subunits of the PP2A family of phosphatases (B’α, B’β, B’’α and B’’β) that interact with PIF3 (Figure 1A and Supplemental Figure S1). B’α and B’β belong to the B’ subfamily, and B’’α and B’’β are from B’’ subfamily with high sequence similarity (Supplemental Figure S2A-B). However, there is low sequence similarity among all four subunits (Supplemental Figure S2C), and the B’ and B’’ form distinct clades in a phylogenetic tree (Farkas et al., 2007; Wang et al., 2016). Although, the substrates for B’α and B’β are known (Tang et al., 2011), no substrates were reported for B’’α and B’’β in *Arabidopsis*. Because PP2A is a heterotrimeric complex in plants, we examined if PIF3 also interacts with the A and C subunits. However, no interaction was detected between PIF3 and PP2A A and C subunits (Supplemental Figure S3).

**Figure 1:**
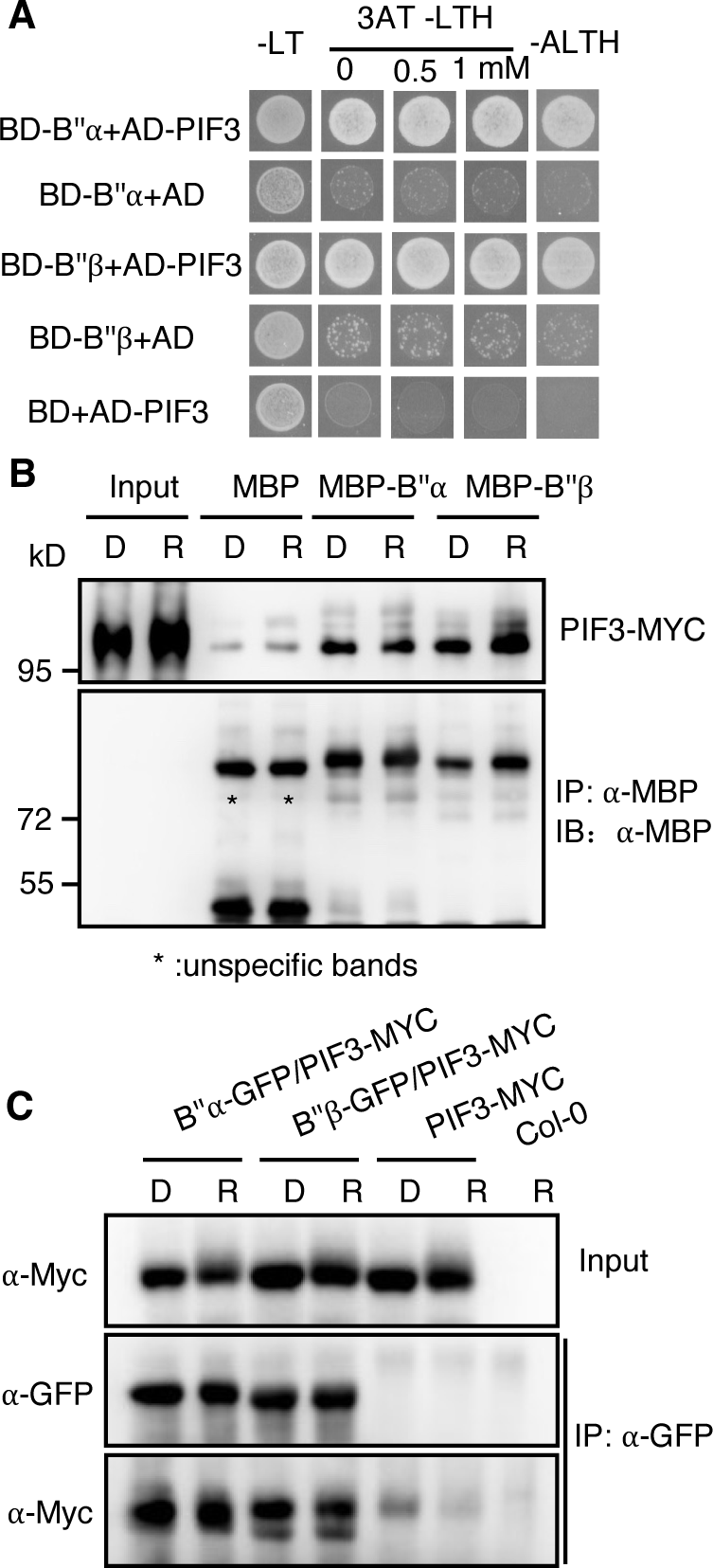
PIF3 interacts with PP2A B’’α and B’’β *in vitro* and *in vivo*. **A.** Yeast two-hybrid assays of the interaction between full-length PIF3, and PP2A B’’α and B’’β subunits. The B’’α- and B’’β-GAL4-DNA binding domain (BD-B’’α and BD-B’’β) fusion was co-expressed with GAL4-activation domain (AD-) fused to full-length PIF3 or AD by itself as a negative control. Yeast cells were grown on selective media lacking histidine, supplemented with an increasing concentration of the histidine biosynthesis inhibitor 3-amino triazole (3-AT). **B.** Semi-*in vivo* pull-down assay shows the interaction between PIF3-MYC and MBP-B’’α and MBP-B’’β. MBP-B’’α and MBP-B’’β proteins were incubated with the extracts from four-day dark-grown seedlings of PIF3-MYC transgenic line (Dark or red-light treated) and then were pull down by MBP beads. Finally, PIF3-MYC signals were detected by α-Myc. MBP only as a negative control. Inputs from Dark and red-light treated extracts as positive controls. **C.** *In vivo* co-immunoprecipitation assay shows that PIF3-MYC interacts with B’’α-GFP and B’’β-GFP in response to red light or in the dark condition. Four-day dark-grown seedlings of 35s:B’’α-GFP/PIF3-MYC, 35s:B’’β-GFP/PIF3-MYC, PIF3-MYC and Col-0 were used. PIF3-MYC and Col-0 were as negative controls. All the seedlings were treated with 100 μM Bortezomib for 4 hours in darkness. One batch was kept in dark condition and the other batch was treated with red light. α-GFP antibody was used to immunoprecipitate the B’’α-GFP and B’’β-GFP and the α-MYC antibody was used to detect PIF3-MYC protein.

To verify the interactions between PIF3 and B subunits, semi-*in vivo* pull-down assay was performed. B’’α and B’’β fused with maltose-binding protein (MBP) were used to incubate with the extracts from dark and light-exposed seedlings of 35S promoter driven *PIF3-MYC* overexpression line and used the MBP beads to pull down the MBP protein and then detected the PIF3-MYC signals. As shown in Figure 1B, PIF3-MYC signals were detected from the dark or red-light treated conditions. Furthermore, co-immunoprecipitation (co-IP) was conducted by using stable transgenic plants (*35S:B’’α-GFP/35S:PIF3-MYC* and *35S:B’’β-GFP/35S:PIF3-MYC*) to verify their interactions. As expected, B’’α-GFP and B’’β-GFP show strong interactions with PIF3-MYC *in vivo* independent of light (Figure 1C). We also performed yeast-two-hybrid interaction assays, semi-*in vivo* pull-down and co-IP experiments and found that PIF3 interacts with B’α and B’β (Supplemental Figure S1A-C). In summary, these results suggest that PP2A B’α, B’β, B’’α and B’’β interact with PIF3 *in vitro* and *in vivo*.

Since major PIFs (PIF1, PIF3, PIF4, and PIF5) share high similarity within protein sequence (Castillon et al., 2007; Lee and Choi, 2017), we tested whether other PIFs (PIF1, PIF4 and PIF5) could interact with PP2A B’’α and B’’β. Interestingly, PP2A B’’α and B’’β could interact with PIF1, PIF4 and PIF5 in yeast and in semi-*in vivo* pull-down experiments (Supplemental Figure S4A-D), suggesting that B’’α and B’’β interact with four major PIFs and might regulate their abundance.

### PP2A B subunits promote hypocotyl elongation under red-light

To examine whether PP2A B’’α and B’’β function in photomorphogenesis, we first isolated T-DNA insertion lines of *b’’α* (SALK_135978) and *b’’β* (SALK_151964) and created overexpression lines using 35S promoter driving *B’’α* and *B’’β* fused with GFP at the C-terminus. Semi-RT PCR assays confirmed that the expression of *B’’α* and *B’’β* are undetected in these mutants, respectively (Supplemental Figure S5). Since PIF3 is involved in regulating hypocotyl elongation under red light (Kim et al., 2003; Monte et al., 2004), we examined the hypocotyl phenotype of the *pp2ab’’α* and *b’’β* single mutants and the overexpression lines under dark and red light conditions. *pp2ab’’α* and *b’’β* exhibited shorter hypocotyl lengths specifically under red-light conditions compared to Col-0 wild type (WT) (Supplemental Figure S6). Conversely, overexpression lines of B’’α (*B’’α-OX*) and B’’β (*B’’β-OX*) exhibited longer hypocotyls than that of WT in red light condition but not in the dark condition (Figure 2E-H). To eliminate the redundancy, we generated the double mutant of *b’’α* and *b’’β* and then analyzed the hypocotyl phenotype. *pp2ab’’αβ* double mutant (or just called *b’’αβ*) shows shorter hypocotyl length compared to WT (Figure 2A-D), like the *b’’β* single mutant under red light conditions (Supplemental Figure S6). However, the hypocotyl length of *b’’αβ* was similar to the WT under darkness, suggesting that the role of PP2A B’’α and B’’β is red light specific (Figure 2A and 2C; Supplemental Figure S6). Intriguingly, the hypocotyl length of *b’’αβ* double mutant was not as short as that of *pif3* mutant, suggesting that other phosphatases may be involved in this process.

**Figure 2:**
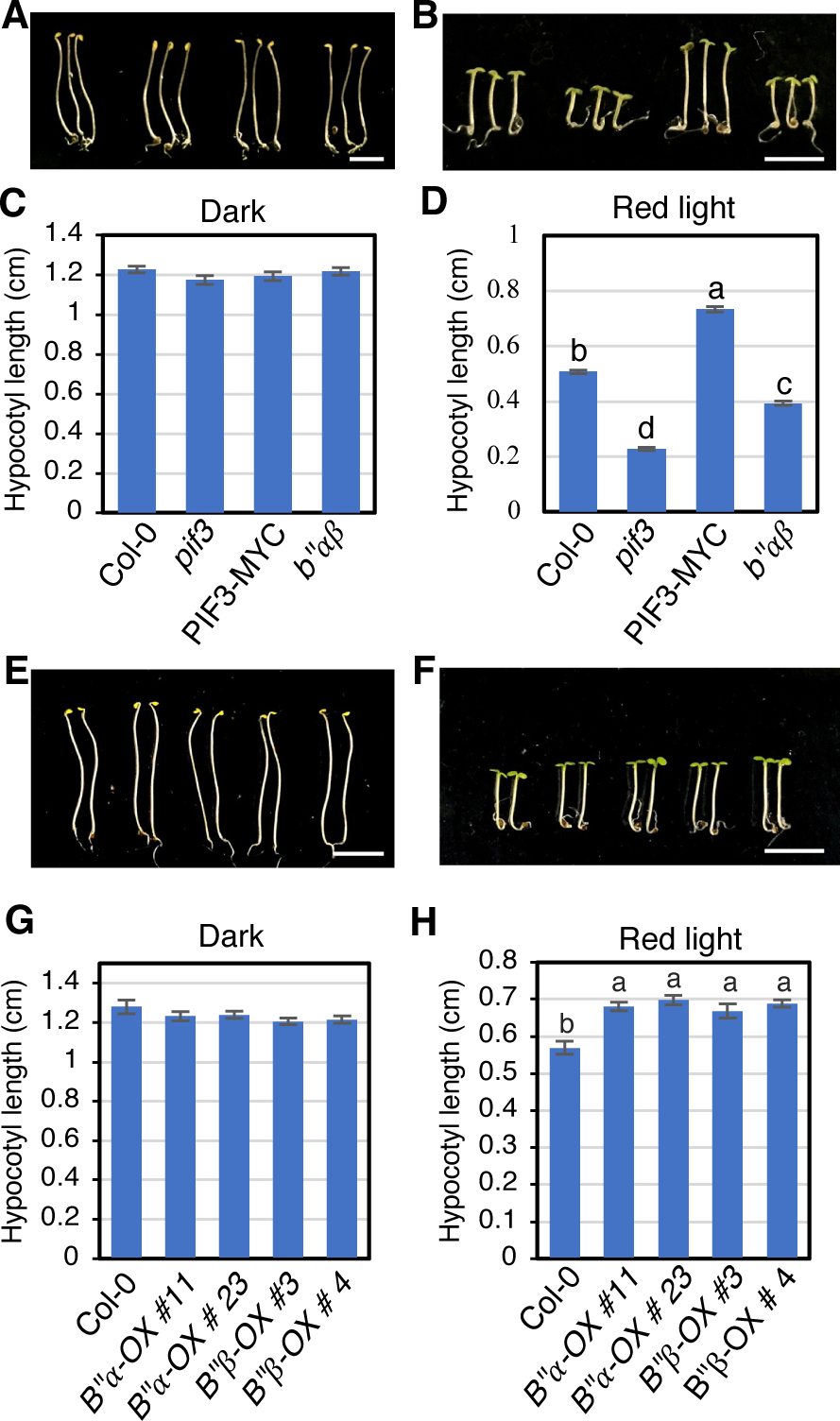
PP2A B’’*α* and B’’*β* promote hypocotyl elongation in red-light condition. **A and B.** Photographs showing the seedling phenotypes of *pp2ab’’αβ* grown in darkness (A) and red-light (8 μmol/m^2^s) condition (B) respectively for 4 days. The seedling order in image from left to right: Col-0, *pif3,* PIF3-MYC and *pp2ab’’αβ*. Scale bar in A and B: 5 mm. **C and D.** Bar graphs show the hypocotyl lengths of seedlings shown in A and B. (*n* ≥ 24). Error bars represent SE. One-way ANOVA was performed. Statistically significant differences are indicated by different lowercase letters (*P* < 0.05). **E and F.** Photographs showing the seedling phenotypes of B*’’α* overexpression lines (*B’’α* #11 and #23) and B*’’β* overexpression lines (*B’’β* #3 and #4) grown in darkness (E) and red-light condition (8 μmol/m^2^s) respectively for 4 days. The seedling order in image from left to right: Col-0, B*’’α* overexpression lines #11and #23, B*’’β* overexpression lines #3 and #4. Scale bar in E and F: 5 mm. **G and H.** Bar graphs show the hypocotyl lengths of seedlings shown in E and F. (*n* ≥10). Error bars represent SE. One-way ANOVA was performed. Statistically significant differences are indicated by different lowercase letters (*P* < 0.05).

To test if *B’α* and *B’β* also function in photomorphogenesis, like *B’’α* and *B’’β*, we examined the hypocotyl phenotype of *b’αβ* double mutant and *B’α-OX* line. Similar to *b’’αβ*, *b’αβ* exhibits short hypocotyl while the *B’α-OX* line shows long hypocotyl under red-light condition compared to wild type, respectively (Supplemental Figure S7A-H). Interestingly, *b’αβ* shows an even stronger hypocotyl phenotype than that of *b’’αβ*. However, *b’αβ* and *B’α-OX* line exhibit a slightly shorter and longer hypocotyl phenotype compared to WT in the dark condition, respectively (Supplemental Figure S7A-H), suggesting that B’α and B’β might also function in the dark condition as has been shown previously (Tang et al., 2011). To test if these four B subunits are functioning redundantly in regulating hypocotyl elongation, we crossed *b’’αβ* and *b’αβ* and generated the *pp2ab’’αβ/b’αβ* quadruple mutant. The hypocotyl phenotype of *pp2ab’’αβ/b’αβ* is as strong as that of *b’αβ* (Supplemental Figure S7A-D). Taken together, these data indicate that B subunits promote hypocotyl elongation in red-light conditions.

To test if PP2A A subunits are involved in regulating photomorphogenesis, we generated two independent lines of the 35S promoter driven overexpression lines of RCN1 (*RCN1-OX)* and A3 (*A3-OX*) with C-terminal GFP tag in WT background and examined the hypocotyl elongation phenotype under the same condition. As expected, both *RCN1-OX* and *A3-OX* display longer hypocotyls in red light condition compared to WT (Supplemental Figure S8A-D), indicating that PP2A A subunits also contribute to hypocotyl elongation. In summary, these data indicate that PP2A acts as a negative regulator of photomorphogenesis under red light condition.

### PIFs and B’’α and B’’β function in the same genetic pathway to regulate hypocotyl elongation

Because B’α, B’β, B’’α, B’’β and PIF3 regulate hypocotyl elongation, we tested if B’’α, B’’β and PIFs function in the same genetic pathway to regulate hypocotyl elongation. To answer this question, we generated *cr-pif3* and *cr-pif3 b’’αβ* triple mutants by using CRISPR-Cas9 to mutate *PIF3* in the WT and *b’’αβ* backgrounds. Western blot results show no expression of PIF3 protein in these *pif3* CRISPR lines (Supplemental Figure S9E). There was no difference in hypocotyl length among genotypes including WT and *cr-pif3b’’αβ* triple mutants under dark condition (Supplemental Figure S9A and C). However, under red-light condition, all the *pif3* CRISPR lines in WT background show short hypocotyls compared to WT (Supplemental Figure S9B and D). The triple mutant shows even shorter hypocotyls than the *pif3* CRISPR lines (Supplemental Figure S9B and To further explore the genetic relation between PIFs and B’’α and B’’β, we crossed *pifQ* with *b’’αβ* and generated the *pifQ/b’’αβ* sextuple mutant and examined the hypocotyl lengths under dark and red-light conditions. Results show that *pifQ* and *pifQ/b’’αβ* sextuple mutant exhibit the same hypocotyl length in the dark and red-light conditions (Figure 3A-D), suggesting *pifQ* is epistatic to *b’’αβ*. To further confirm the genetic relationship between *PIF3* and *B’’β*, *35S:B’’β-GFP/pif3* was generated and hypocotyl length was measured. Results show that the long hypocotyl phenotype of *B’’β-OX* is largely eliminated in the *B’’β-OX/pif3* background under red light condition (Figure 3E to 3H), indicating that *B’’β-OX* phenotype is PIF3-dependent. Western blot analysis confirmed that B’’β-GFP protein level did not alter in Col-0 and *pif3* background (Supplemental Figure S10A and B), indicating that PIF3 mutation caused the short hypocotyl length of *B’’β-OX/pif3*, not the protein level of B’’β-GFP. These data demonstrate that PIFs and B’’α, B’’β function in the same genetic pathway to regulate hypocotyl elongation under red light.

**Figure 3:**
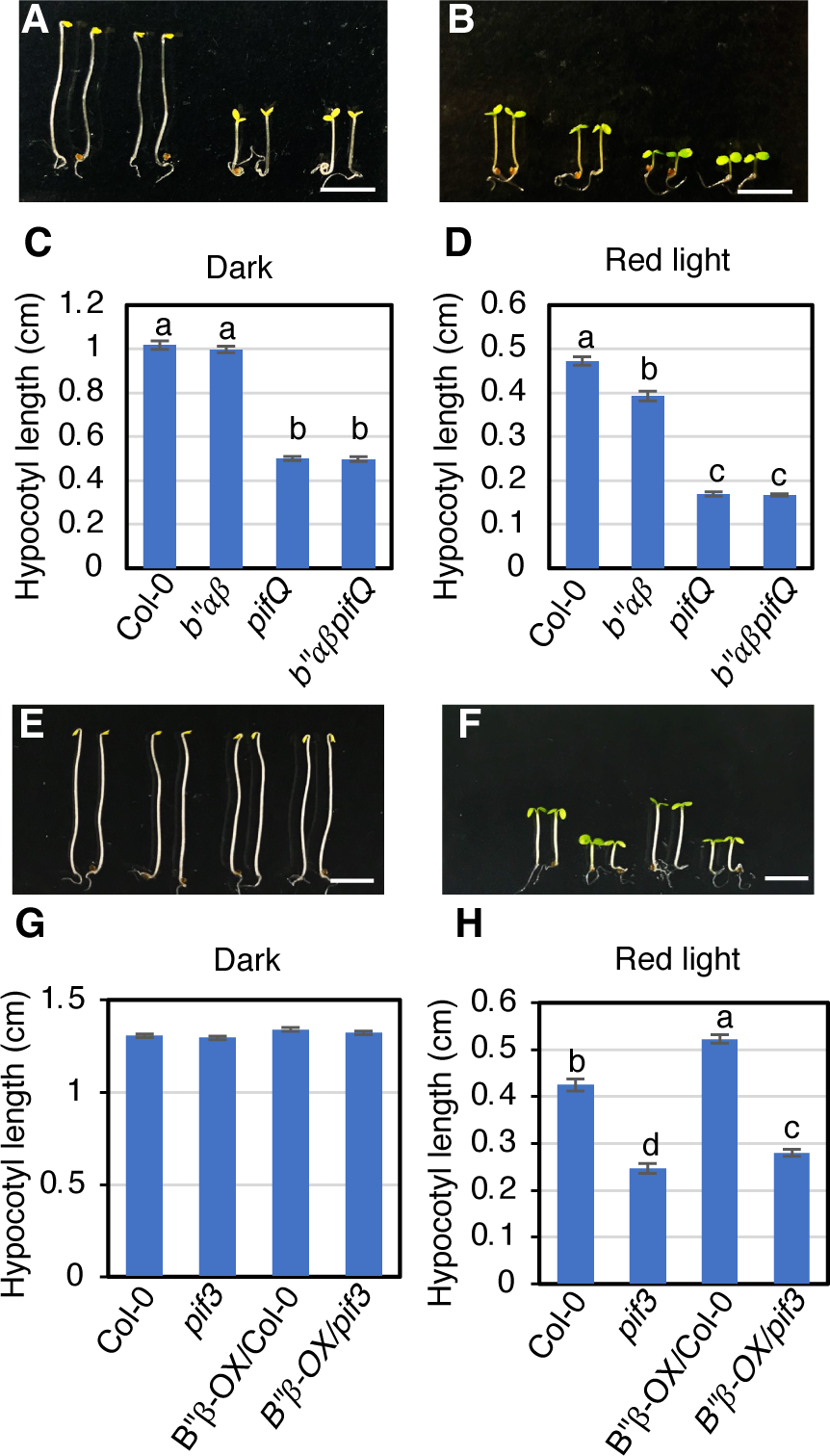
PP2A B*’’α* and B*’’β* and PIFs act in the same genetic pathway to regulate hypocotyl elongation in *Arabidopsis*. **A and B.** Photographs showing the seedling phenotypes grown in darkness (A) and red-light condition (B, 8 μmol/m^2^s) respectively for 4 days. The seedling order in image from left to right: Col-0, *pp2ab’’αβ, pifQ, b’’αβpifQ.* Scale bar in A and B: 5 mm. **C and D.** Bar graphs show the hypocotyl lengths of seedlings shown in A and B. (*n* ≥ 30). Error bars represent SE. One-way ANOVA was performed. Statistically significant differences are indicated by different lowercase letters (*P* < 0.05). **E and F.** Photographs showing the seedling phenotypes grown in darkness (A) and red-light condition (B, 8 μmol/m^2^s) respectively for 4 days. The seedling order in image from left to right: Col-0, *pif3, B’’β-OX/Col-0, B’’β-OX/pif3.* Scale bar in A and B: 5 mm. **G and H.** Bar graphs show the hypocotyl lengths of seedlings shown in E and F. (*n* ≥ 20). Error bars represent SE. One-way ANOVA was performed. Statistically significant differences are indicated by different lowercase letters (*P* < 0.05).

To provide additional genetic evidence, we also used the CRISPR-Cas9 method to mutate *B’’α* and *B’’β* simultaneously in the *35S:PIF3-MYC* background. We identified one *35S:PIF3-MYC/cr-b’’αβ #32* line. Sequencing result revealed that this line contains 1 bp deletion in *B’’α* and 1bp insertion in *B’’β* (Supplemental Figure S11A), which cause frameshift and lead to early termination (Supplemental Figure S11B and C). Phenotypic analysis showed that *35S:PIF3-MYC*/*cr-b’’αβ #32* exhibits shorter hypocotyls than *35S:PIF3-MYC* under red light conditions (Supplemental Figure S12), suggesting that the *B’’α* and *B’’β* promote PIF3 function possibly by inhibiting red-light induced degradation of PIF3.

To substantiate these genetic data, we used a pharmacological approach and treated WT, *pif3*, *35S:PIF3-MYC*, *b’’αβ* double mutant and *B’’α-OX* and *B’’β-OX* seedlings with cantharidin, a widely used inhibitor of protein phosphatase types 1 (PP1) and PP2A (Honkanen, 1993) under red light and checked the hypocotyl length. As shown in Supplemental Figure S13A, with the increasing concentration of cantharidin, the hypocotyl elongation was gradually inhibited among all the genotypes. Fifteen μM Cantharidin can abolish long hypocotyl phenotype of *35S:PIF3-MYC*, *B’’α-OX* and *B’’β-OX* under red-light condition. When the relative hypocotyl length change was calculated by using hypocotyl length values from 15 μM Cantharidin condition divided by values of DMSO control, *pif3* (around 6% reduction) is hyposensitive to Cantharidin and *b’’αβ* double mutant (around 33% reduction) is less sensitive to Cantharidin treatment compared to WT (around 43% reduction). However, *35S:PIF3-MYC* (around 60% reduction), *B’’α-OX* (around 52% reduction) and *B’’β-OX* (around 50% reduction) are hypersensitive to Cantharidin (Supplemental Figure S13B). These data suggest that PP2A and/or possibly PP1 activity is critical for the longer hypocotyl phenotype of *35S:PIF3-MYC*, *B’’α-OX* and *B’’β-OX* under red light.

### PP2A dephosphorylates and stabilizes PIF3 from red-light induced degradation

During dark to red-light transition, PIF3 is first phosphorylated and then degraded by the 26S proteasome pathway (Park et al., 2004; Al-Sady et al., 2006). We hypothesized that B subunits may have a role in PIF3 dephosphorylation since B’α, B’β, B’’α and B’’β belong to a phosphatase family. Therefore, we examined native PIF3 level in WT and *b’’αβ/b’αβ* quadruple mutant using 4-day-old dark-grown seedlings and dark-grown seedlings exposed to red light over time. PIF3 was gradually degraded in both WT and the *b’’αβ/b’αβ* quadruple mutant after red-light treatment. However, the degradation rate of native PIF3 in *b’’αβ/b’αβ* quadruple mutant was faster than that in WT background (Figure 4A, B). When we performed the PIF3 degradation assay by using WT and *b’’αβ* double mutant, PIF3 also showed faster degradation in the *b’’αβ* double mutant compared to WT (Supplemental Figure S14A and B). These data suggest that the B subunits inhibit the red-light-induced degradation of PIF3.

**Figure 4:**
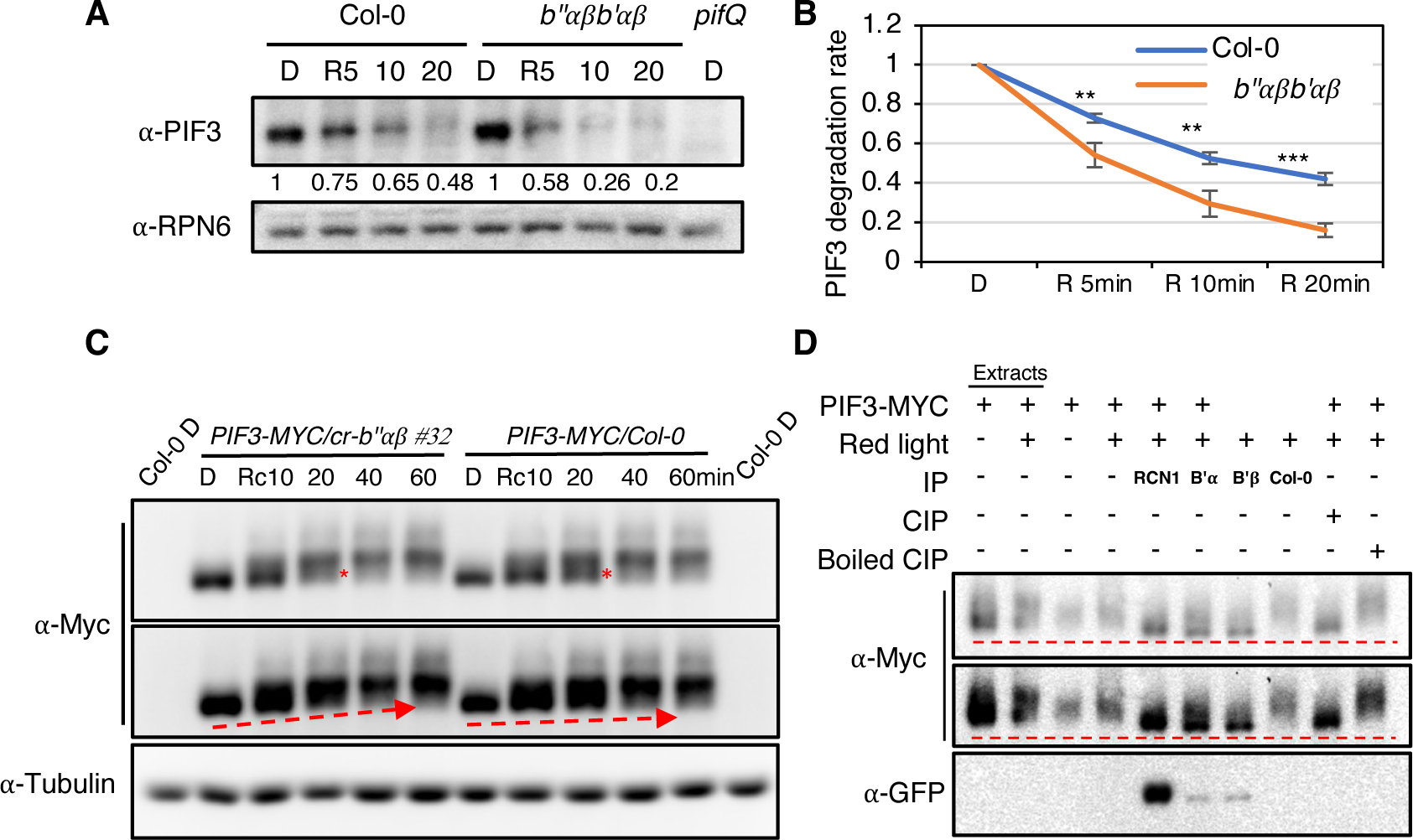
PP2A controls PIF3 level under red light by dephosphorylation. A. Immunoblots showing the light-induced degradation of native PIF3 in the *pp2ab’’αβb’αβ* mutant compared to wild type. Four-day-old dark-grown seedlings were either kept in darkness or exposed to red light (20 μmol/m^2^s) for the duration indicated before being sampled for protein extraction. RPN6 blot as loading control. The numbers show the native PIF3 protein abundance after calibrating with RPN6 bands. The assay was repeated independently twice with similar results. **B.** Line graphs show native PIF3 degradation rate after red light exposure in wild type and *pp2ab’’αβb’αβ* mutant background based on three independent blots. ** P<0.01 and *** P<0.001, based on Student’s t-test. Error bars represent SE (n=3). **C.** Immunoblots show the light-induced phosphorylation of PIF3-MYC in the *cr-b’’αβ #32* mutant and wild type. Four-day-old dark-grown seedlings were treated with 100 μM Bortezomib for 4 hours and then either kept in darkness or exposed to red light (20 μmol/m^2^s) for the duration indicated. Tubulin blot shows loading control. Red asterisks indicate the shift difference between *cr-b’’αβ #32* mutant and wild type. The red dash arrows show the phosphorylation trend between *cr-b’’αβ #32* mutant and wild type. **D.** PP2A dephosphorylates PIF3 *in vitro*. Dephosphorylation assay was performed by using immunoprecipitated PIF3-MYC and PP2A proteins from PIF3-MYC plants and RCN1-GFP, YFP-B’*α* and YFP-B’*β* transgenic plants, respectively. PIF3-MYC proteins from four dark-grown seedlings of PIF3-MYC, treated with 100 μM Bortezomib for 4 hours in darkness and exposed to red-light before the immunoprecipitation. RCN1-GFPαYFP-B’*α* and YFP-B’*β* immunoprecipitation (IP) products as PP2A phosphatase incubate with immunoprecipitated PIF3-MYC for 1 hour at 30℃. CIP as positive control. Boiled CIP and IP product from Col-0 as negative controls. CIP and Boiled CIP treatments were performed at 37 ℃ for 1 hour. Western-blot analyses was performed with anti-Myc on SDS-PAGE.

To test if the phosphorylation level of PIF3 is altered in the *b’’αβ* compared to WT, we performed immunoblot analysis of PIF3-MYC levels in *PIF3-MYC/Col-0* and *PIF3-MYC/cr-b’’αβ #32* backgrounds. A time course analysis shows that the phosphorylated form of PIF3-MYC is more abundant in the *PIF3-MYC/cr-b’’αβ #32* background compared to *PIF3-MYC/Col-0* transgenic lines only under red light conditions (Figure 4C). These data suggest that B’’α and B’’β prevent PIF3 degradation by regulating PIF3 phosphorylation status under red light.

To examine if PP2A can directly dephosphorylate PIF3 *in vitro*, we conducted the dephosphorylation assay using immunoprecipitated PIF3-MYC and different PP2A subunits from *35S:PIF3-MYC/Col-0, PP2A 35S:A3-GFP/Col-0, RCN1-GFP/Col-0, B’’β-GFP/Col-0, YFP-B’α/Col-0* and *YFP-B’β/Col-0* transgenic plants, respectively. We used the A3-GFP, RCN1-GFP, B’’β-GFP, YFP-B’α and YFP-B’β immunoprecipitation (IP) products as PP2A phosphatase and performed the incubation with immunoprecipitated PIF3-MYC. As shown in the Figure 4D and Supplemental Figure S14C and S15A, A3-GFP, RCN1-GFP, YFP-B’α and YFP-B’β immunoprecipitated products can directly dephosphorylate PIF3-MYC, similar to the positive control Calf Intestinal Phosphatase (CIP) treatment. However, B’’β-GFP immunoprecipitated products failed to dephosphorylate PIF3-MYC (Supplemental Figure S15A), like the boiled CIP negative control. To investigate the reason for the inability of B’’β-GFP to dephosphorylate PIF3-MYC, we checked whether B’’β-GFP can associate with the PP2A C subunit. As shown in the Supplemental Figure S15B, only a small amount of the C subunit was found to co-precipitate with the B’’β-GFP compared to RCN1-GFP, which suggests that B’’β-GFP exhibits lower binding ability to the C subunit compared with RCN1-GFP. Interestingly, YFP-B’α and YFP-B’β show normal binding to the C subunit compared to B’’β-GFP (Supplemental Figure S16). To test whether the C-terminal GFP fusion is causing reduced association with the C subunit, we generated the *35S:B’β-GFP/Col-0* transgenic plants and examined the PP2A C subunit after performing the immunoprecipitation from YFP-B’β and B’β-GFP plants. In this assay, we used RCN1-GFP, YFP-B’α as positive controls and B’’β-GFP as negative control. The results show that B’β-GFP exhibits lower binding ability compared to YFP-B’β. Quantitative data show that YFP-B’α and YFP-B’β exhibit similar strong binding ability to the C subunit among these subunits tested. However, B’β-GFP only shows 1/3 binding ability to the C subunit compared to YFP-B’β. B’’β-GFP displays the lowest binding ability to the C subunit among of these subunits tested (Supplemental Figure S16). These data suggest that a free C-terminus might be essential for B subunits to bind to the C subunit for PP2A holoenzyme assembly and that B’’β-GFP failed to dephosphorylate PIF3-MYC possibly due to the c-terminal GFP tag.

### PP2A B’’α and B’’β alter gene expression to response to red-light

Red light induces global changes in gene expression to regulate photomorphogenesis (Tepperman et al., 2004; Pfeiffer et al., 2014). To test whether *PP2A B’’α* and *B’’β* can regulate gene expression in response to red light, RNA-seq was conducted by using four-day-old dark-grown seedling of WT and *b’’αβ* mutant, kept in the dark or 1-hour red light (8 μmol·m^-2^·s^-1^) treatment before the samples were collected. The results show that 1359 genes were differentially expressed in the WT and 1387 genes were regulated in *b’’αβ* mutant in response to 1 hour of red-light exposure (Figure 5A; Dataset S1). In addition, 512 genes were *B’’α-* and *B’’β*-dependent, which is around 38% of the DEGs (Dataset S1). Furthermore, 1359 DEGs in WT displayed different patterns in *b’’αβ* as shown in the heatmap analysis (Figure 5B). Gene Ontology (GO) analysis of 512 PP2A B’’α- and B’’β-dependent genes shows an enrichment of the genes involved in response to endogenous, external, osmotic and temperature stimulus and regulation of biological processes and others (Figure 5C). Overall, these data suggest that *PP2A B’’α* and *B’’β* are crucial for transcriptional regulation in photomorphogenesis.

**Figure 5:**
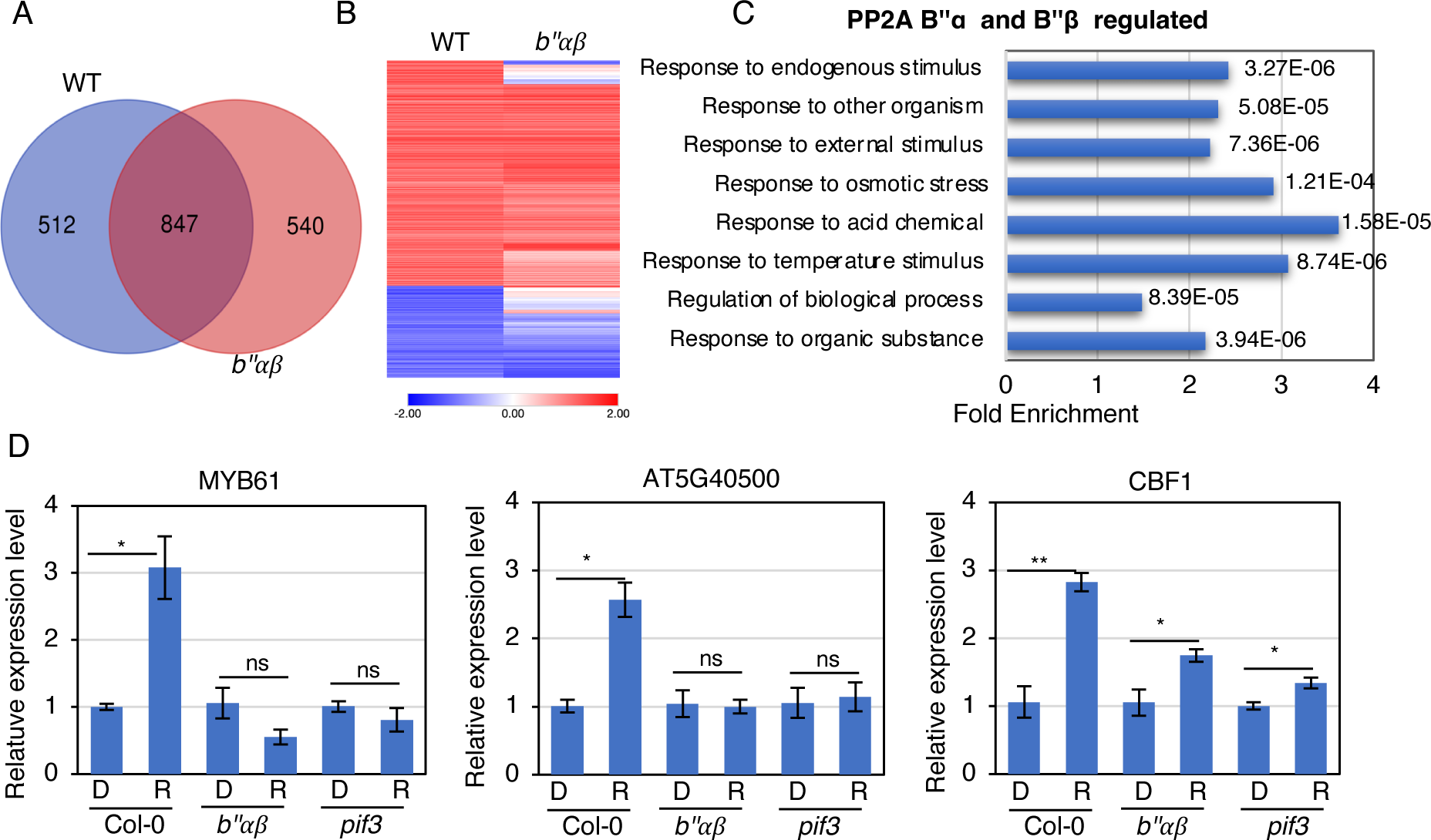
RNA-sequencing revealed unique roles of PP2A B’’α and B’’β in gene expression after red-light exposure. **A.** Venn diagram shows differentially expressed genes (DEGs) in wild-type (WT) vs *pp2ab’’αβ* mutant after red-light exposure. Four-day dark-grown seedlings were exposed to continuous red light (20 μmol/m^2^s) for 1 hour or keep in the darkness and total RNA was extracted from three biological replicates for RNA-seq analyses. **B.** Hierarchical clustering from 1359 DEGs from WT show distinct pattern in *pp2ab’’αβ* mutant after red-light exposure. **C.** Gene Ontology (GO) analysis of PP2A B’’α and B’’β-dependent 512 genes. **D.** RT-qPCR analysis using MYB61, AT5G40500, and CBF1. RT-qPCR samples were from four-day dark-grown seedlings of Col-0, *pp2ab’’αβ* and *pif3* and then either keep in dark condition or exposed to continuous red light (20 μmol/m^2^s) for 1 hour. Three biological repeats were performed. Error bars represent SE (n=3). Relative gene expression levels were normalized using the expression level of ACT2 and the values of those genes in dark condition. Student t-test was performed. * represents *P* < 0.05, * * represents *P* < 0.01.

To confirm the RNA-seq data, we perform qPCR assays to examine the transcript level of several red-light responsive genes, such as *MYB61*, *CBF1* and *AT5G40500* (Figure 5D). Consistent with the RNA-seq data, the transcript levels of these genes were upregulated in response to red light in WT background. However, the transcript levels show either no change or slightly upregulated after red-light treatment in *b’’αβ* and similar trends were observed in *pif3* background. These data suggest that PP2A B’’α and B’’β might modulate gene expression through regulating PIF3 level in response to red-light. In summary, these data indicate that *PP2A B’’α* and *B’’β* play an important role in photomorphogenesis via transcriptional regulation.

### The expression and stability of PP2A subunits are modestly regulated by light

To examine the subcellular localization of the PP2A B’’α and B’’β subunits, we examined the B’’α-GFP and B’’β-GFP localization by using *B’’α-OX* and *B’’β-OX* lines, respectively. The strong GFP signals were found in the nucleus and cytoplasmic area (Supplemental Figure S17). Thus, PP2A B’’ subunits might function in both nucleus and cytoplasm. To test whether the stability of the B’’ subunits is regulated by light, 4-day-old dark-grown seedlings were exposed red light for 1 and 6 hours and the protein level was examined using anti-GFP antibody. Results show that both B’’α and B’’β subunits are modestly stabilized under light conditions (Supplemental Figure S18). We also performed RT-qPCR to test if the expression of any of the *PP2A* subunits is regulated by light. Results show that the expression of *B’α*, *A2* and *C5* is slightly upregulated under red light while the expression of the *C1* subunits is slightly reduced under 6 h of red light (Supplemental Figure S19). Finally, we examined if the expression of the A and C subunits is altered in the *b’’αβ/b’αβ* quadruple mutant background under dark and light conditions. Results show that the expression of *RCN1*, *A2*, *C1*, *C2* and *C3* is not altered in the *b’’αβ/b’αβ* quadruple mutant background compared to WT controls (Supplemental Figure S20). These data suggest that light signals have modest, if any, impact on the expression and stability of the PP2A subunits.

## Discussion

The reversible phosphorylation of proteins especially transcription factors play crucial roles in regulating the physiology of all organisms and has been implicated in almost all signaling pathways (Stark, 2004). An attachment of a phosphate by a kinase and the removal of the phosphate by a phosphatase provide a reversible way of regulating the activity, abundance and/or subcellular localization of a protein. PIFs are a group of basic helix-loop-helix (bHLH) transcription factors and function as negative regulators of photomorphogenesis. In recent years, it has been shown that PIFs are first phosphorylated by multiple kinases and then degraded through the ubiquitin/26S proteasome pathway (Cheng et al., 2021). Our data and those of others show that PIF3 is also dephosphorylated by multiple phosphatases to fine tune photomorphogenesis (Yu et al., 2019).

Several lines of evidence support our conclusion that PP2A regulates photomorphogenesis by dephosphorylating PIF3. First, PP2A B’α, B’β, B’’α and B’’β interact with PIFs *in vitro* and *in vivo*. (Figure 1, Supplemental Figures S1 and S4). Second, *pp2ab’’αβ* and *b’’αβ/b’αβ* seedlings exhibit short hypocotyls in red-light conditions compared to WT in a light-dependent manner (Figure 2A-D, Supplemental Figure S7). Conversely, B’α, B’’α and B’’β overexpression lines show longer hypocotyls compared to WT (Figure 2A-D, Supplemental Figure S7). Third, B’’α and B’’β and PIFs function in the same genetic pathway to regulate hypocotyl elongation (Figure 3, Supplemental Figure S15). Fourth, the light-induced degradation of PIF3 is faster in *b’’αβ/b’αβ* and *b’’αβ* compared to WT background (Figure 4A and B, Supplemental Figure S14A and B). Fifth, the phosphorylation level of PIF3 under red light is higher in the *b’’αβ* background compared to PIF3-MYC transgenic line (Figure 4C). Sixth, PP2A directly dephosphorylates PIF3-MYC *in vitro* (Figure 4D, Supplemental Figures S14C and S15A). Seventh, PP2A B’’α and B’’β regulate gene expression in response to red light (Figure 5). Taken together, these data firmly establish that PP2A functions as a negative regulator of photomorphogenesis.

Previously, two phosphatases have been reported to regulate PIFs phosphorylation status and abundance. The first reported phosphatase to regulate PIFs’ stability is TOPP4, a catalytic subunit of the PP1 family (Yue et al., 2016). TOPP4 inhibits the red light-induced ubiquitination and degradation of PIF5 during photomorphogenesis in *Arabidopsis*. *topp4-1* mutant displayed short hypocotyls and expanded cotyledon compared to wild type under red light. Further protein interaction assays and phosphorylation studies demonstrate that TOPP4 interacts directly with and dephosphorylates PIF5. However, TOPP4 was isolated as a dwarf mutant suggesting a more general function in regulating plant growth and development (Qin et al., 2014). The other phosphatases to regulate PIFs are Fypp1 and Fypp2, two genes encoding the catalytic subunits of protein phosphatases 6 (PP6) (Yu et al., 2019). Fypp1 and Fypp2 directly interact with PIF3 and PIF4 in *Arabidopsis.* However, two evidence suggest that Fypp1 and Fypp2 mainly function in the dark condition. First, *fypp1* and *fypp2* mutants show short hypocotyls in the dark condition compared to WT, but slightly longer than *pifQ*. Second, PIF3 and PIF4 proteins exhibited mobility shifts in *fypp1fypp2* mutants due to their hyperphosphorylation in the dark conditions compared with that in the Col-0 background. Thus, PP2A B subunits are different from previously reported phosphatases that regulate PIFs’ stability and phosphorylation status only under light conditions. Based on our data, B’’α and B’’β are two PP2A regulatory subunit genes that specifically function in the red-light condition because *b’’αβ* or *B’’α-OX* and *B’’β-OX* lines only display hypocotyl phenotypes compared to WT specifically under red-light conditions (Figure 2, Supplemental Figures S6A-D and S7A-D).

Although our genetic and biochemical data support the conclusion that PP2A regulates photomorphogenesis by dephosphorylating PIF3, the change in phosphorylation status of PIF3 in the *cr-b’’αβ #32* background is rather modest (Figure 4C). PIF3 has been shown to be phosphorylated in a large number of serine/threonine residues (Ni et al., 2013). It is possible that PP2A B’ and B’’ subunits dephosphorylate only a subset of these phosphorylation sites. This is consistent with our Cantharidin treatment data, suggesting other phosphatases including PP1 family may also participate in the PIF3 dephosphorylation process. Identifying the exact dephosphorylation sites might help understand the role of PP2A B’ and B’’ subunits in this process.

Although, PP2A B subunits regulate photomorphogenesis, the hypersensitive phenotypes of the *b’αβ*, *b’’αβ* and *b’’αβ/b’αβ* are rather weak under red light conditions. There are more than 17 B subunits in the PP2A family in *Arabidopsis* (Farkas et al., 2007; Booker and DeLong, 2016), suggesting that the gene redundancy may contribute to the weak hypocotyl phenotypes (Figure 2A-D, Supplemental Figures S6 and S7). It is highly possible that other B subunits also are involved in this process. Our pharmacological data support this conclusion where *pp2ab’’αβ* is hyposensitive to Cantharidin, but *B’’α-OX* and *B’’β-OX* lines are hypersensitive to Cantharidin (Supplemental Figure S13), meaning there are other subunits and/or classes of phosphatases involved in this process. Interestingly, *pif3* is hyposensitive to Cantharidin but *PIF3-MYC* is hypersensitive to Cantharidin (Supplemental Figure S13), which suggests PIF3 function required PP2A or PP1 family of phosphatases. Overall, these data suggest that phosphatases play crucial roles in regulating PIF abundance and activity to modulate photomorphogenesis.

One unexpected finding from our co-IP assays is that a free c-terminus is essential for B subunits to associate with C subunits *in vivo*. Based on our IP results, only B subunits with a free c-terminus (YFP-B’α or YFP-B’β) show stronger binding to the C subunit (Supplemental Figures S15B and S16). However, B’β-GFP or B’’β-GFP exhibit lower binding ability to the C subunit compared to YFP-B’α or YFP-B’β (Supplemental Figure S16). These data suggest that B subunits promote hypocotyl elongation relying on the binding ability to the C subunit of PP2A. B’’β-GFP only shows longer hypocotyl phenotype when the expression level is very high. Even though B’’β-GFP expression level is much higher than YFP-B’α (Supplemental Figure S16), *YFP-B’α* (*B’α-OX*) exhibits much longer hypocotyl compared to wild type (Supplemental Figure S7), suggesting a free C-terminus promotes association with the C subunit and enhance the role of B subunits in regulating photomorphogenesis.

In summary, our data show that PP2A B subunits directly interact with PIFs and dephosphorylate PIF3 to inhibit its degradation to fine-tune photomorphogenesis. PP2A is a new group of phosphatases that regulate PIFs phosphorylation and stability in the photomorphogenesis process along with other phosphatases. Thus, multiple kinases and phosphatases regulate PIF abundance and activity to regulate photomorphogenesis (Figure 6).

**Figure 6.**
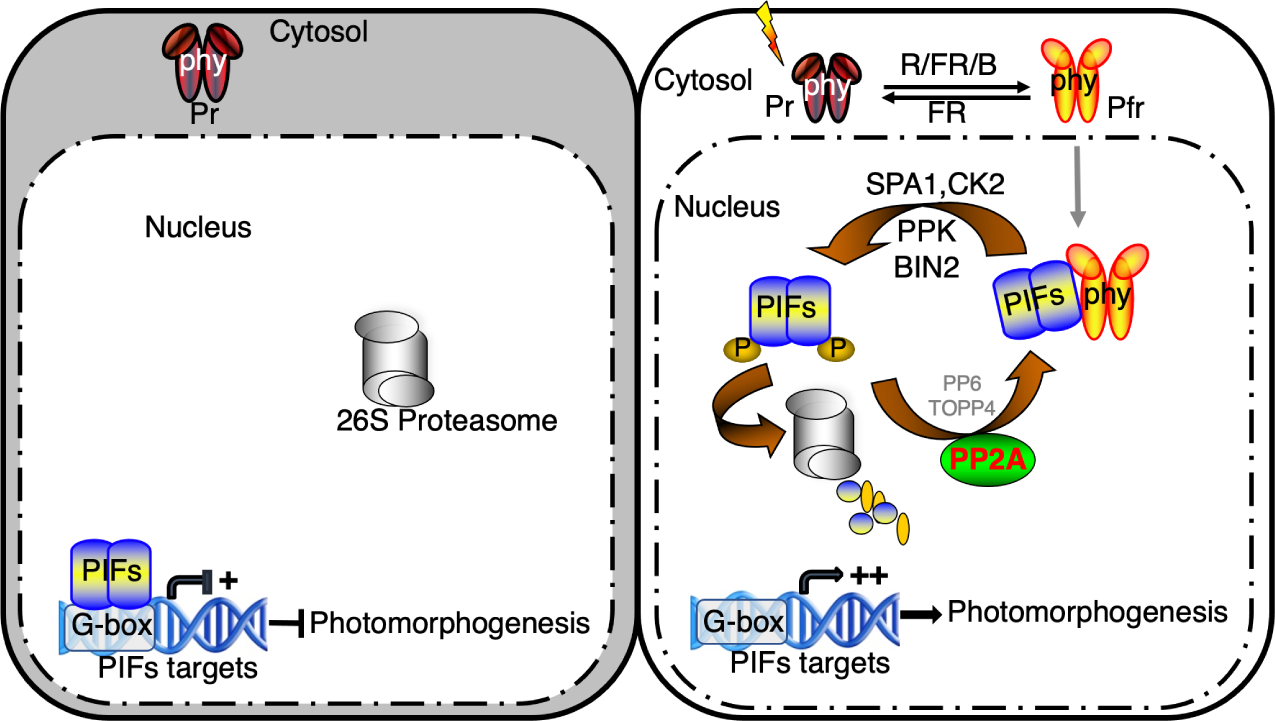
Model of phytochrome signaling pathway. Left panel: In the dark, phytochromes are in an inactive Pr form and stay in the cytosol. And the nuclear-localized PHYTOCHROME INTERACTING FACTORs (PIFs) can form homo or heterodimers or tetramers to bind to the promoter region of their target genes to repress their expression and prevent photomorphogenesis. Right panel: Upon light exposure, phytochromes convert from Pr form to active Pfr form and translocate into the nucleus. In the nucleus, the interaction between phytochromes and PIFs triggers the rapid phosphorylation of PIFs by several kinases (SPA1, CK2, PPKs, BIN2). The phosphorylated PIFs will be degraded by the 26S proteasome pathway. The degradation of the PIFs promotes light-regulated gene expression and photomorphogenesis. Conversely, PP2A with other phosphatases (TOPP4 and PP6) can dephosphorylate PIFs to inhibit their degradation to fine-tune photomorphogenesis.

## Materials and methods

### Plant growth conditions and hypocotyl phenotype analysis

All the seeds are of *Arabidopsis thaliana* in Columbia-0 (Col-0) background. The T-DNA insertion mutants used in this paper: *b’’α* (SALK_135978), *b’’β* (SALK_151964) and *b’αβ* (Tang et al., 2011). Seeds were surface sterilized in 1% bleach solution with 0.3% SDS for 10 min followed by five quick rinses with sterile water and then plated on Murashige and Skoog (MS) basal salts without sucrose. Seeds were stratified at 4°C in the dark for 3 days followed by white light exposure for 3 hours to promote germination and then either kept in the dark (22°C) or continuous red light (light intensity and time of exposure indicated in each figure) for 4 days for Western blot analysis or hypocotyl elongation analysis. Hypocotyl length was measured using ImageJ software and analyzed by using one-way ANOVA or students t-test. Statistical differences are indicated by different letters or asterisks (P < 0.05).

### Construction of vectors and generation of transgenic plants

To generate *B’’α-OX*, *B’’β-OX*, *RCN1-OX*, *A3-OX* transgenic plants, B’’α, B’’β, RCN1, A3 coding sequence were amplified using primers listed in Table S1 and first subcloned into pENTR vector (Thermofischer Scientific, Cat# K240020), respectively. pENTR-B’α and pENTR-B’β vectors have been previously described (Wang et al., 2016). Then pENTR-B’’α, B’’β, B’β, RCN1, A3 were recombined to pB7FWG2 gateway binary vector with 35S promoter (Karimi et al., 2005) and c-GFP tag by using LR Clonase II (Thermofischer Scientific, Cat# 11791020). Then pB7FWG2 destination vectors with genes were transformed into Col-0 background, respectively and transformants were selected in the presence of Basta antibiotic. Homozygous lines were selected from T3 generation with detectable GFP signals. *YFP-B’α* and *YFP-B’β* transgenic plants have been previously described (Tang et al., 2011).

To generate *crispr-b’’αb’’β*, two sets of target sites were selected from close to N-terminus of *B’’α* and *B’’β* genes, respectively. pHEE401E was chosen as a destination vector (Wang et al., 2015). Then pHEE401E-B’’αB’’β was transformed into *PIF3-MYC* background and transformants were selected in the presence of Hygromycin B and Kanamycin antibiotics. Homozygous lines were selected by sequencing.

### Protein purification from E. coli

For MBP-B’α, B’β, B’’α and B’’β, LR reactions were performed between pENTR-B genes and pVP13 vector with MBP tag (Jeon et al., 2005). Each plasmid was transformed into BL21(DE3) cells. Protein expression was induced under 16 °C for 12 h with 0.1 mM IPTG. Extraction buffer (50 mM Tris, pH 7.5, 150 mM NaCl, 1 mM EDTA, 0.1% Tween20, 0.25 mM DTT, 1X protease inhibitor cocktail, 1 mM PMSF) was added to cell pellet and vertex to resuspend cell. Sonication was performed to break the cells and the extracts were cleared by centrifugation at 20000 × g for 15 min. The supernatants were incubated with amylose resin (NEB, Cat#, E8021S) for one hour in the dark at 4°C. Amylose resin was washed with extraction buffer three times for 10 mins each time. The MBP protein was still bound to resin for the following *in vitro* pull-down assays.

### Protein interaction assays

For semi-*in vitro* pull-down assays, four-day old dark-grown seedlings of *35S:PIF3-MYC* were treated with 100 μM Bortezomib for 4 hours in darkness. One batch was ground in liquid nitrogen. Another batch was exposed to 20 μmol·m^-2^·s^-1^ red light for 10s and then kept in dark until 10 min before being ground in liquid nitrogen. Total protein was solubilized in extraction buffer (50 mM Tris, pH 7.5, 150 mM NaCl, 1 mM EDTA, 0.1% Tween-20, 0.25 mM DTT, 1 × protease inhibitor cocktail (Sigma-Aldrich, Cat# P9599), 1 mM PMSF). The extracts were cleared by centrifugation at 20000 × g for 15 min. The supernatants were incubated with beads bound with MBP-B’’α, MBP-B’’β, and MBP only as a control, respectively for one hour in the dark at 4 °C. Beads were washed three times, 10 mins each with extraction buffer. Beads were then boiled with 2X SDS sample buffer and the supernatants were separated on an SDS-PAGE gel. Anti-MYC antibody was used to detect PIF3-MYC protein.

For the *in vivo* co-immunoprecipitation assays, four-day old dark-grown *35S:B’’α-GFP/PIF3-MYC*, *35S:B’’β-GFP/PIF3-MYC*, *35S:PIF3-MYC* and Col-0 seedlings were used. *35S:PIF3-MYC* and Col-0 were used as controls. The seedlings were treated with 100 μM Bortezomib for 4 hours in darkness. One batch was ground in liquid nitrogen. Another batch was exposed to 20 μmol·m^-2^·s^-1^ red light for 10s and then kept in dark until 10 min before being ground in liquid nitrogen. Total protein was solubilized in extraction buffer (50 mM Tris, pH 7.5, 150 mM NaCl, 1 mM EDTA, 0.1% Tween20, 0.25 mM DTT, 1X protease inhibitor cocktail, 1 mM PMSF). The extracts were cleared by centrifugation at 20000 × g for 15 min. The supernatants were incubated with Dynabeads bound with anti-GFP (Abcam, Cat. # ab6556, 20 μL/μg) for one hour in the dark at 4°C. Dynabeads were washed with extraction buffer three times for 10 mins each time. Beads were heated to 65°C for 15min with 2X SDS sample buffer and separated on an 8% SDS-PAGE gel. Anti-Myc antibody was used to detect PIF3-MYC protein. Anti-GFP antibody was used to detect B’’α-GFP and B’’β-GFP proteins.

For yeast two-hybrid assay, AD plasmids (AD-PIF1, AD-PIF4, AD-PIF5), and BD plasmids (BD-B’’α, BD-B’’β) were transformed into the AH109 yeast cell simultaneously and then selected on a solid medium lacking Leu and Trp amino acids (-LT). Only successfully transformed yeast cells survive on -LT medium. Then these yeast cells were plated on a solid medium lacking Leu, Trp, and His amino acids (-LTH). The synthesis of His can be activated when B’’α and B’’β interact with PIF1, PIF4, and PIF5, and then the yeast can survive on the -LTH medium. To avoid false-positive results, 3-amino-1,2,4-triazole (3-AT) were added to the -LTH medium to inhibit the self-activation of the His synthesis.

### PIF3 degradation assay

To observe native PIF3 degradation in Col-0 and *pp2ab’’αβ*, four-day old dark-grown seedlings were used. Seedlings were exposed to red light (20 μmol·m^-2^·s^-1^) for 0, 5, 10, and 20 min before being collected, and total protein was extracted in extraction buffer (8M urea, 50 mM Tris-Cl, pH 7.5, 1X protease inhibitor cocktail). Anti-PIF3 primary antibody from Agrisera (Cat# AS16 3954) was used and the secondary antibody also is from Agrisera (Cat# AS09605). For the loading control, anti-RPT5 (Enzo Life Sciences, Cat # BML-PW8770-0025), anti-RPN6 (Agrisera, Cat# AS152832A) or Coomassie Blue (CBB) staining was used.

### PIF3 dephosphorylation assay

#### IP for PIF3-MYC

Four-day old dark-grown seedlings of PIF3-MYC were treated with 100 μM Bortezomib for 4 hours in the dark and then was exposed to 20 μmol·m^-2^·s^-1^ red light for 10s and then kept in dark until 10 min before being ground in liquid nitrogen. The sample was ground in the buffer (50 mM Tris-Cl (pH=7.5), 150 mM NaCl, 1% Triton-X 100, 1mM PMSF, 100 μM Bortezomib, 1× protease inhibitor cocktail, 25 mM β-glycerophosphate, 10 mM NaF, and 2 mM Na orthovanadate), and immunoprecipitated using anti-Myc (Sigma, Cat. # C3956) antibody pre-bound to Dynabeads. The immunoprecipitated PIF3-MYC was eluted with 0.1M Glycine (pH=2.0) from Dynabeads and then neutralized with 1M Tri-Cl (pH=8.0) (v/v=2/1). Eluted PIF3-MYC was aliquoted into different microtubes as substrate

#### IP for PP2A

Four-day old dark-grown seedlings of RCN1-GFP, A3-GFP and *b’’αβ* (as negative control) were ground in the buffer (50 mM Tris-Cl (pH=7.5), 150 mM NaCl, 0.1% NP-40, 1mM PMSF, 100 μM Bortezomib, 1 × protease inhibitor cocktail, 5mM CaCl_2_). Samples were immunoprecipitated using anti-GFP (Abcam, Cat. # ab290) antibody pre-bound to Dynabeads. IP products were washed once by wash buffer (50 Tris-Cl (pH=7.5), 150 mM NaCl, 0.1% NP-40, 5mM CaCl_2_).

#### Dephosphorylation

Mix the IP products from RCN1-GFP, A3-GFP and *b’’αβ* with PIF3-MYC substrate, respectively, in the buffer (50 mM Tris-Cl (pH=7.5), 150 mM NaCl, 10mM MgCl_2_, 1mM MnCl_2_, 1mM DTT, 1 × protease inhibitor cocktail, 5mM CaCl_2_), incubate in 30°C shaker for 1 hour. Also mix Quick CIP (NEB, Cat. #M0525) or boiled Quick CIP with PIF3-MYC substrate, respectively, in the buffer (50 mM Tris-Cl (pH=7.5), 100 mM NaCl, 10mM MgCl_2_, 1mM DTT, 1× protease inhibitor cocktail), incubate at 37 °C for 1 hour. Western blot was then performed by using 7% gel, and anti-Myc was used to detect PIF3-MYC signals.

### RNA extraction, cDNA synthesis, and qRT-PCR

Four-day-old dark-grown seedlings were used with four independent biological replicates (n=4). The seeds were surface sterilized and plated on the MS media without sucrose, cold stratified for 3 days before treated for 3 hours of white light. After 4 days, seedlings were either kept at dark or exposed to red-light (20 μmol·m^-2^·s^-1^) for 1 hour or 6 hours. Total RNA was isolated using the plant total RNA kit (Sigma, Cat. # STRN250). For cDNA synthesis, 2 μg of total RNA was used for reverse transcription with M-MLV Reverse Transcriptase (Thermofischer Scientific, Cat. # 28025013). SYBR Green PCR master mix (Thermofischer Scientific, Cat. # 4368577) and gene-specific oligonucleotides were used to conduct qPCR analyses using primers shown in Table S1. Finally, relative transcription level was calculated using 2^-ΔΔCT^ using ACT7 normalization.

### RNA-seq analyses

Col-0 (WT) and *pp2ab’’αβ* seedlings were used for mRNA sequence analysis. The seeds were surface sterilized and plated on the MS media without sucrose, cold stratified for 3 days before treated for 3 hours of white light and 21 h of dark incubation to induce germination. After the initial 24 h, seeds were further treated with far-red light (10 μmol·m^-2^·s^-1^) for 5 min to inactivate phytochrome activity. After an additional 3 days in the dark, seedlings were either kept in dark (dark samples) or treated with 1 h of continuous red light (red samples) (20 μmol·m^-2^·s^-1^). Total RNA was isolated using the plant total RNA kit (Sigma, Cat. # STRN250). The 3’Tag-Seq method was employed for RNA-seq analysis in this study (Lohman et al., 2016). FastQC was used for examining raw read quality (www.bioinformatics.babraham.ac.uk/projects/fastqc/). HISAT2 was used to align raw reads to the Arabidopsis genome (Kim et al., 2019). The annotation of the Arabidopsis genome was from TAIR10 (www.arabidopsis.org/). Read count data were obtained using HTseq (Anders et al., 2015) (htseq.readthedocs.io/en/master/). The EdgeR was used to identify the differentially expressed genes in WT/ *pp2ab’’αβ* (Robinson et al., 2010). Cutoff and adjusted P value (FDR) for the differential gene expression was setup with ≥2-fold and ≤0.05 respectively. Heatmap was generated by using Morpheus (https://software.broadinstitute.org/morpheus/) and Venn diagrams were generated using the website (http://bioinformatics.psb.ugent.be/webtools/Venn/). For the heatmap analysis, we used the hierarchical clustering with one minus cosine similarity metric combined with the average linkage method. Also, GO enrichment analyses were performed using (http://geneontology.org). GO bar graphs were generated based on the result of the significantly enriched terms with the lowest P value and FDR (≤0.05) in GO terms. Raw data and processed data for RNAseq in Col-0 and *b’’αβ* can be accessed from the Gene Expression Omnibus database under accession number GSE174428.

## Acknowledgements

We thank Dr. Inyup Paik for helpful discussion and suggestions, members of the Huq laboratory for critical reading of the manuscript. The authors acknowledge the Texas Advanced Computing Center (TACC) at The University of Texas at Austin for providing High Performance Computing, visualization, and database resources that have contributed to the research results reported in this paper.

## Author contributions

E. H. and X.C. conceived the study and designed the experiments. X.C. carried out the experiments. S.L. and X.C analyzed the RNA-seq data. W.T. and Y.S. helped prepare the PP2A genetic materials and Y2H vectors. X.C. and E.H. wrote the article. S.L. Y.S. and W.T. commented and edited the manuscript.

## Data availability

RNA sequencing data were deposited into the Gene Expression Omnibus database (accession number GSE174428). *Arabidopsis* mutants and transgenic lines, as well as plasmids and antibodies generated during the current study are available from the corresponding author upon reasonable request.

---

**Supplemental Figure S1:**
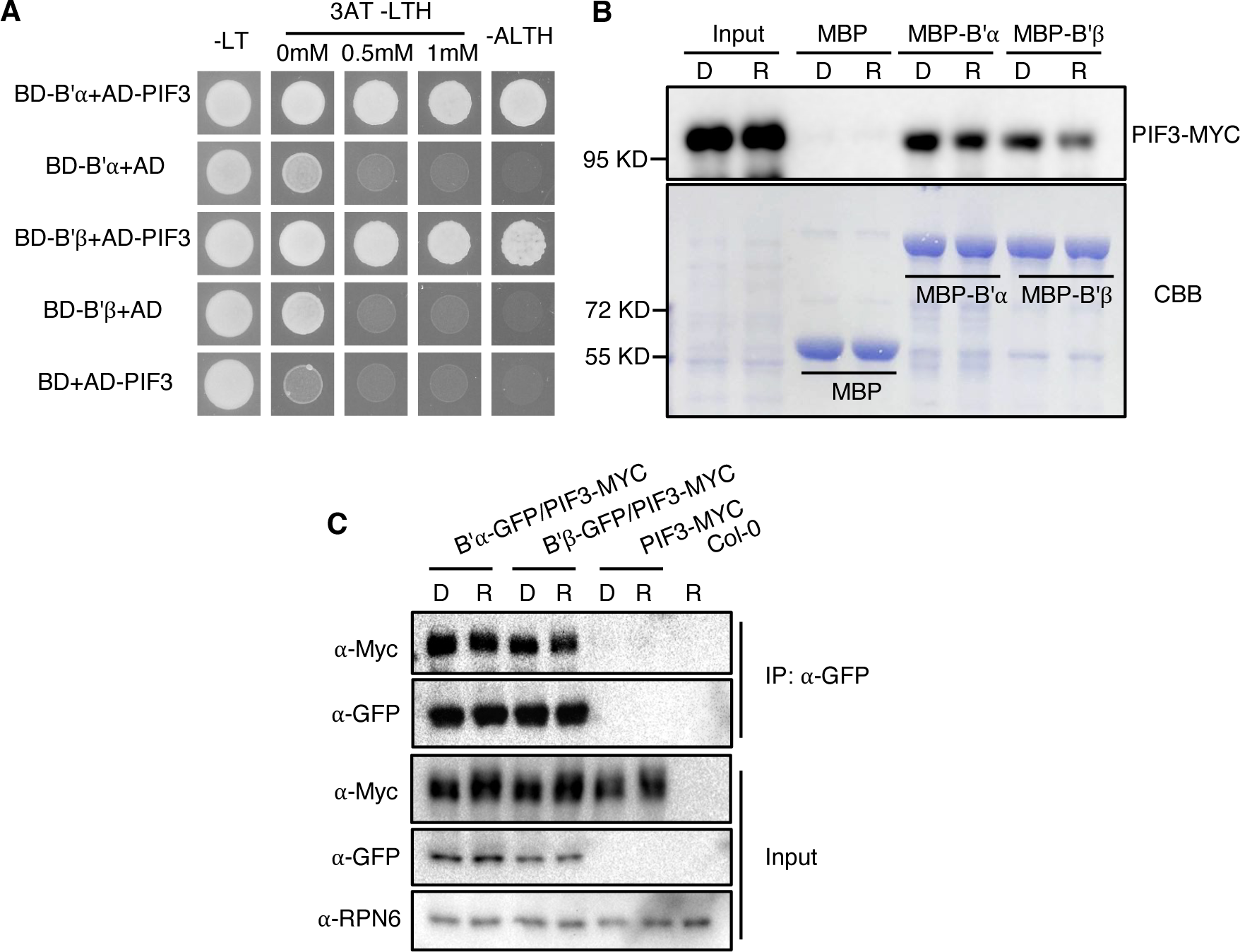
PIF3 interacts with PP2A B’α and B’β in vivo and in vitro. **A.** Yeast two-hybrid assays of the interaction between full-length PIF3, and PP2A B’α and B’β. The B’α- and B’β-GAL4-DNA binding domain (BD-B’α and BD-B’β) fusion was co-expressed with GAL4-activation domain (AD-) fused to full-length PIF3 or AD by itself as a negative control. Yeast cells were grown on selective media lacking histidine, supplemented with an increasing concentration of the histidine biosynthesis inhibitor 3-amino triazole (3-AT). **B.** Semi-in vitro pull-down assay shows the interaction between PIF3-MYC and MBP-B’α and MBP-B’β. MBP-B’α and MBP-B’β proteins were incubated with the extracts from four-day dark-grown seedlings of PIF3-MYC transgenic line (Dark or red-light treated) and then were pull down by MBP beads. Finally, PIF3-MYC signals were detected by α-Myc. MBP only as a negative control. Inputs from Dark and red-light treated extracts as positive controls. 10 % gel was used in this assay. **C.** *In vivo* co-immunoprecipitation assay shows that PIF3-MYC interacts with B’α -GFP and B’β-GFP in response to red light or in the dark condition. Four-day dark-grown seedlings of 35s::B’α-GFP/PIF3-MYC, 35s::B’β-GFP/PIF3-MYC, PIF3-MYC and Col-0 were used. PIF3-MYC and Col-0 were as negative controls. All the seedlings were treated with 100 μM Bortezomib for 4 hours in darkness. One batch was kept in dark condition and the other batch was treated with red light. α-GFP antibody was used to immunoprecipitate the B’α-GFP and B’β-GFP and the α-MYC antibody was used to detect PIF3-MYC protein. RPN6 blot as loading control.

**Supplemental Figure S2:**
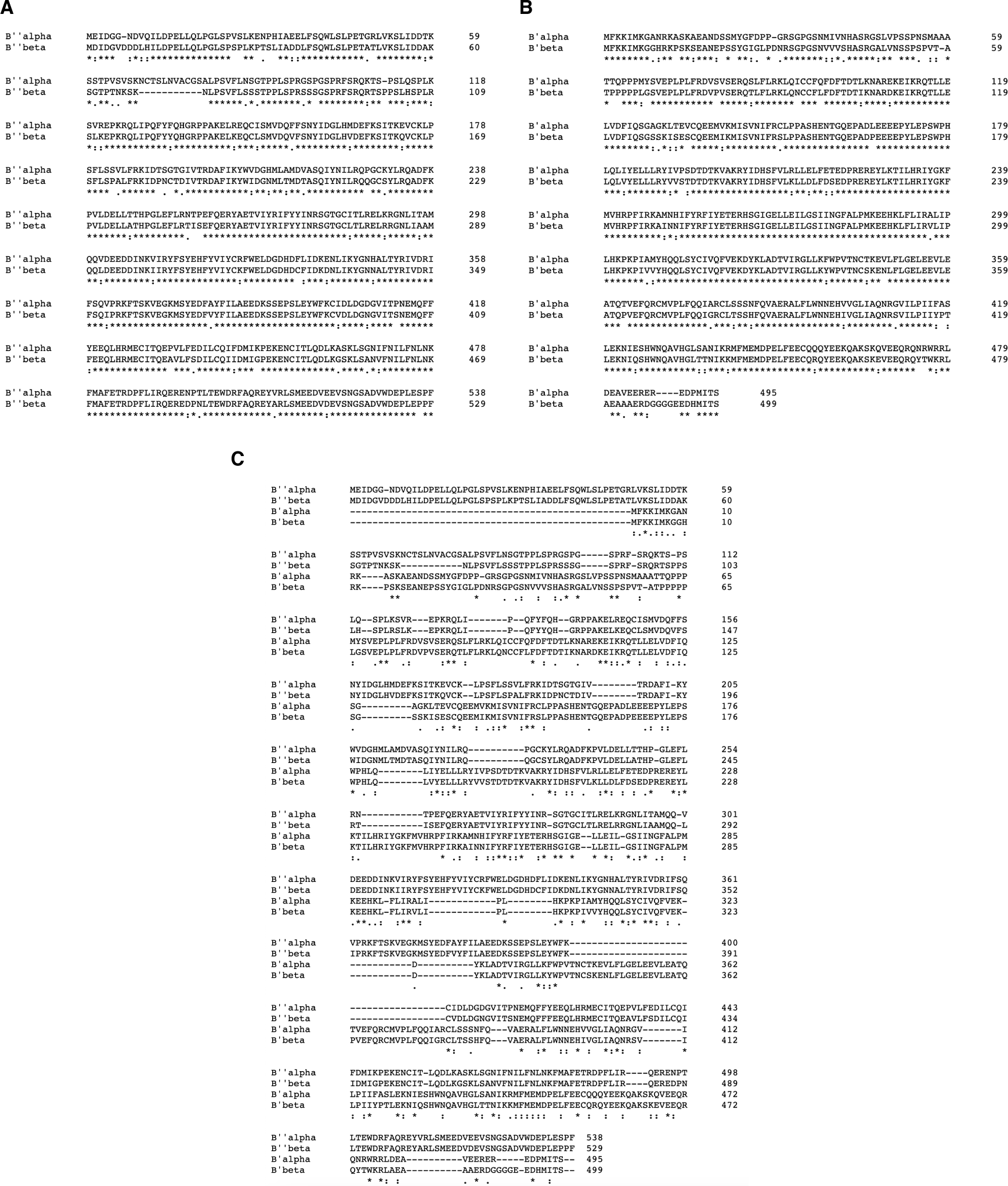
Amino acid sequence alignment among PP2A B subunits. Amino acid sequence alignment between B’’α and B’’β (A), B’α and B’β (B) and all 4 subunits (C). Alignments were performed using Clustal Omega online tool (https://www.ebi.ac.uk/Tools/msa/clustalo/).

**Supplemental Figure S3:**
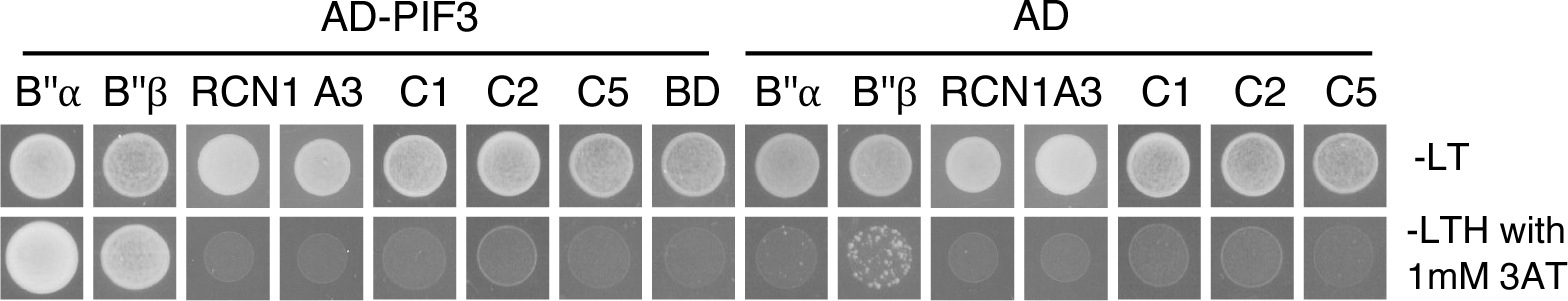
PIF3 doesn’t interact with PP2A A subunits and catalytic (C) subunits in yeast. Yeast two-hybrid assays showing interaction of PIF3 with PP2A B’’α and B’’β, but not with PP2A A subunits (RCN1 and A3) and catalytic subunits (C1, C2 and C5). The B’α- and B’β-, A subunits (RCN1- and A3-) and catalytic subunits (C1-, C2- and C5-)-GAL4-DNA binding domain fusions were co-expressed with GAL4-activation domain (AD-) fused to full-length PIF3 or AD by itself as a negative control. Yeast cells were grown on selective media lacking histidine, supplemented with 1 mM of the histidine biosynthesis inhibitor 3-amino triazole (3-AT).

**Supplemental Figure S4:**
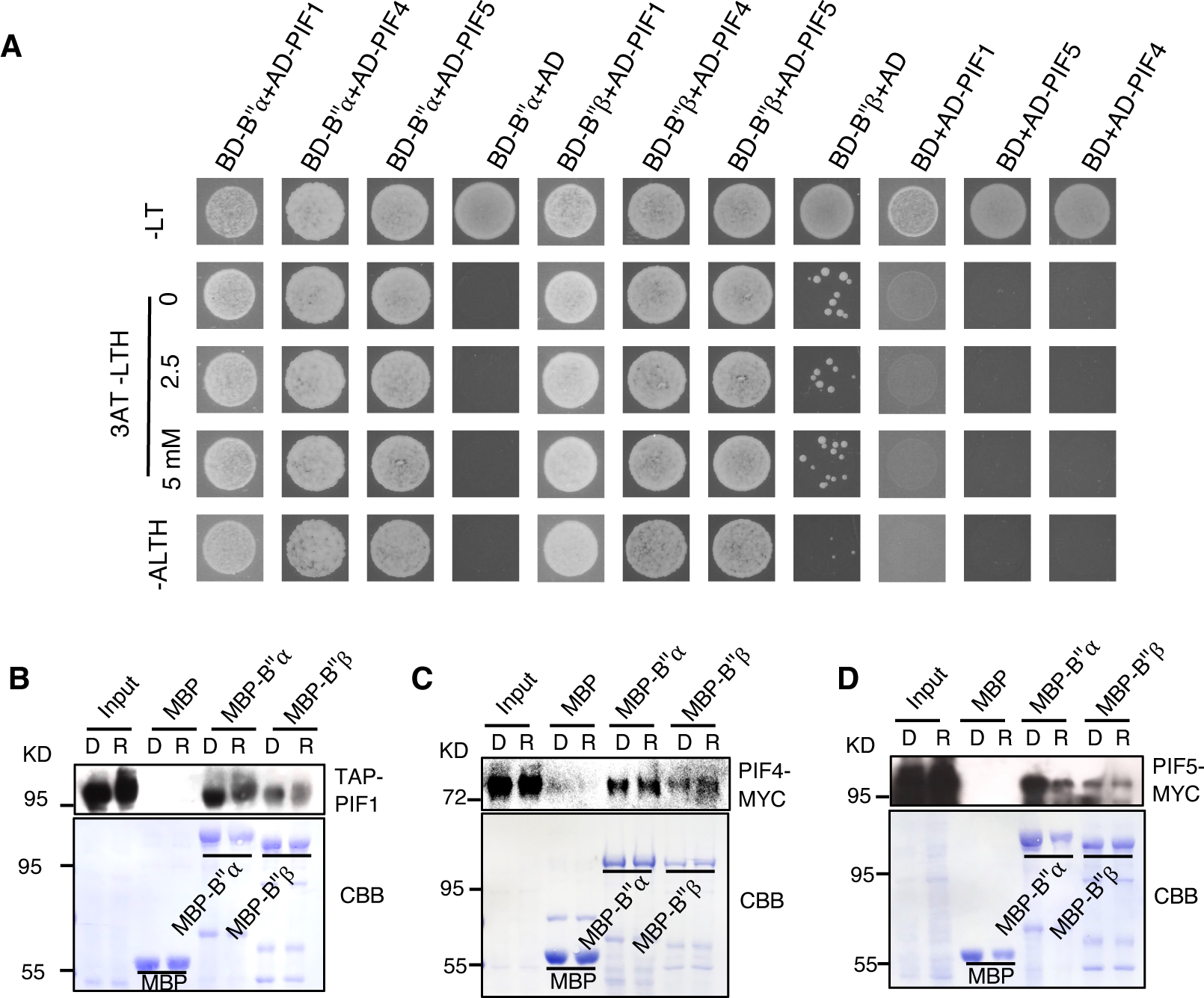
PIF1,4 and 5 interact with PP2A B’’*α* and B’’*β*. **A.** Yeast two-hybrid assays of the interaction between full-length PIF1, PIF4, PIF5 and PP2A B’’*α* and B’’*β*. The B’’α- and B’’β-GAL4-DNA binding domain (BD-B’’α and BD-B’’β) fusion was co-expressed with GAL4-activation domain (AD-) fused to full-length PIF1, PIF4, PIF5 or AD by itself as a negative control. Yeast cells were grown on selective media lacking histidine, supplemented with an increasing concentration of the histidine biosynthesis inhibitor 3-amino triazole (3-AT). **B-D.** Semi-in vitro pull-down assay shows the interactions between TAP-PIF1, PIF4-MYC, PIF5-MYC and MBP-B’’*α* and MBP-B’’*β*. MBP-B’’α and MBP-B’’β proteins were incubated with the extracts from four-day dark-grown seedlings of TAP-PIF1, PIF4-MYC and PIF5-MYC transgenic lines (Dark or red-light treated) and then were pull down by MBP beads. Finally, TAP-PIF1, PIF4-MYC and PIF5-MYC signals were detected by α-Myc antibody. MBP only as a negative control. Inputs from Dark and red-light treated extracts as positive controls

**Supplemental Figure S5:**
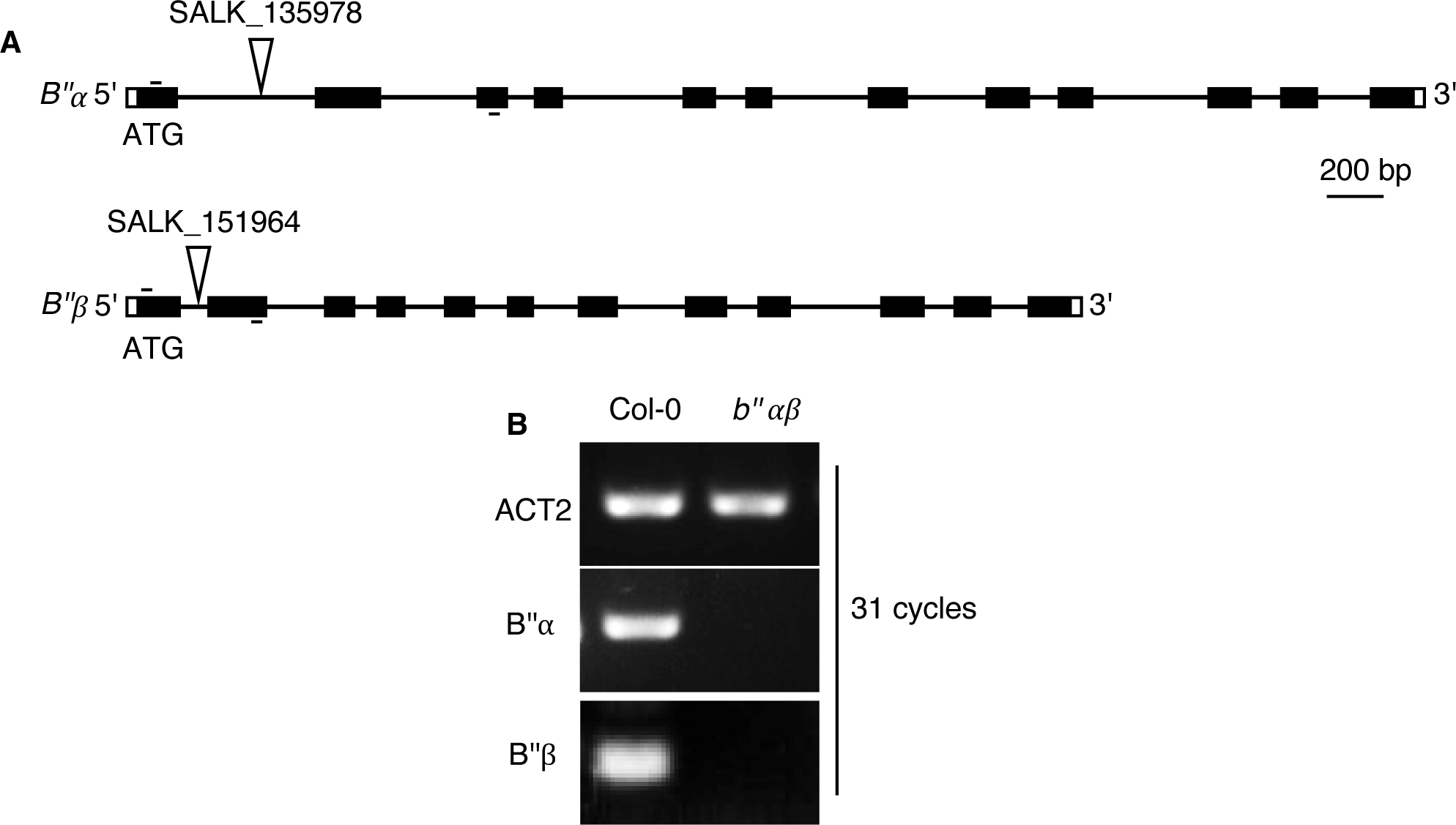
**A.** The structure of the *PP2AB’’α* and *B’’β* genes showing the location of T-DNA insertions. Closed boxes indicate exons; open boxes represent untranslated regions; solid lines indicate introns. T-DNA insertion sites are indicated by triangles. The small black lines on the gene structures show the primer positions. **B.** RT-PCR analysis indicates the *PP2AB’’α* and *B’’β* transcripts are not detectable in the *pp2ab’’αβ* T-DNA mutant.

**Supplemental Figure S6:**
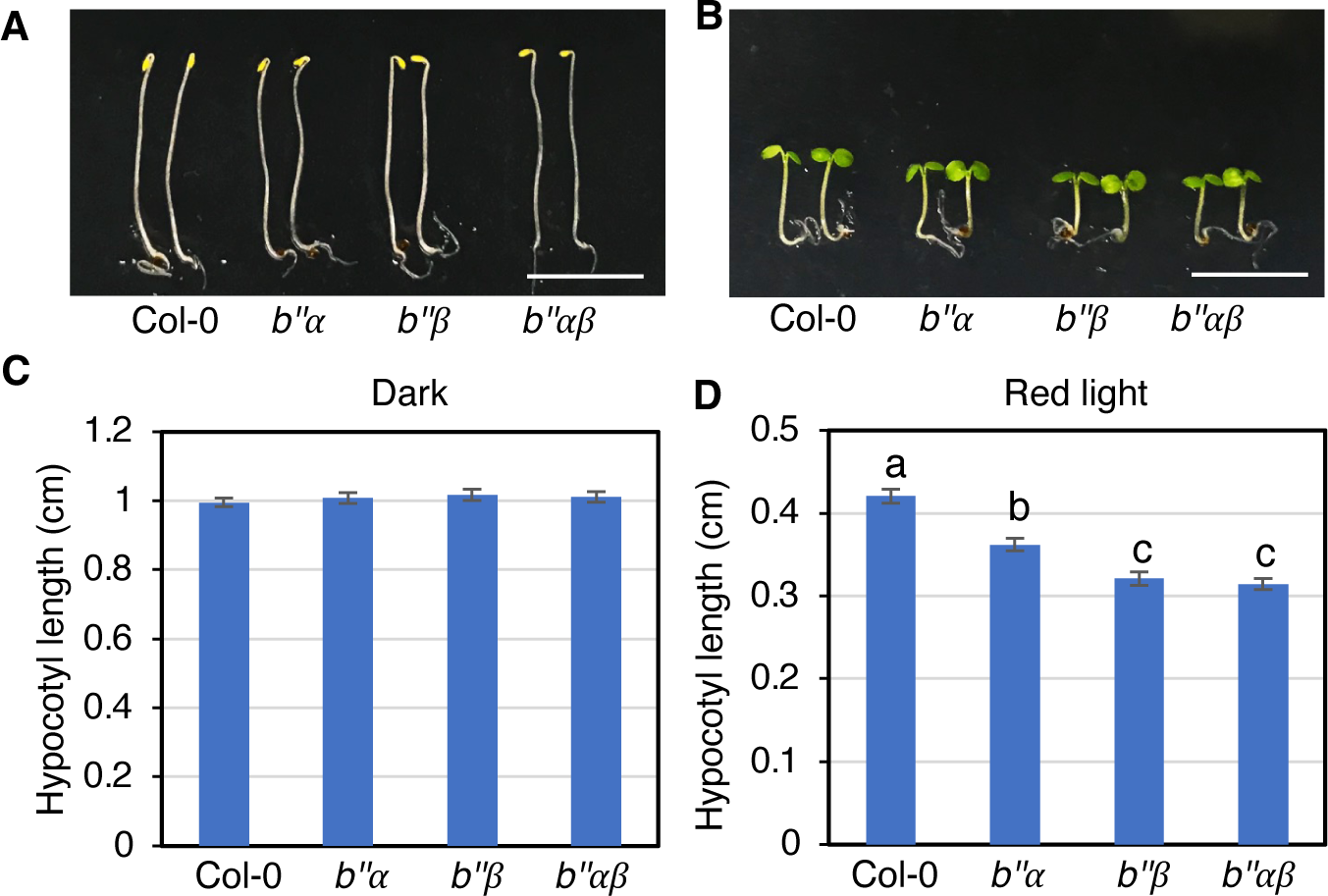
*pp2ab’’α* and *b’’β* seedlings show short hypocotyls in red-light condition. **A and B.** Photographs showing the seedling phenotypes of of *pp2ab’’α* and *pp2ab’’β* grown in darkness (A) and red-light (8 μmol/m^2^s) condition (B) respectively for 4 days. The seedling order in image from left to right: Col-0, *pp2ab’’α, pp2ab’’β* and *pp2ab’’αβ*. Scale bar in A and B: 5 mm. **C and D.** Bar graphs show the hypocotyl lengths of seedlings shown in A and B. (*n* ≥ 28). Error bars represent SE. One-way ANOVA was performed. Statistically significant differences are indicated by different lowercase letters (*P* < 0.05).

**Supplemental Figure S7:**
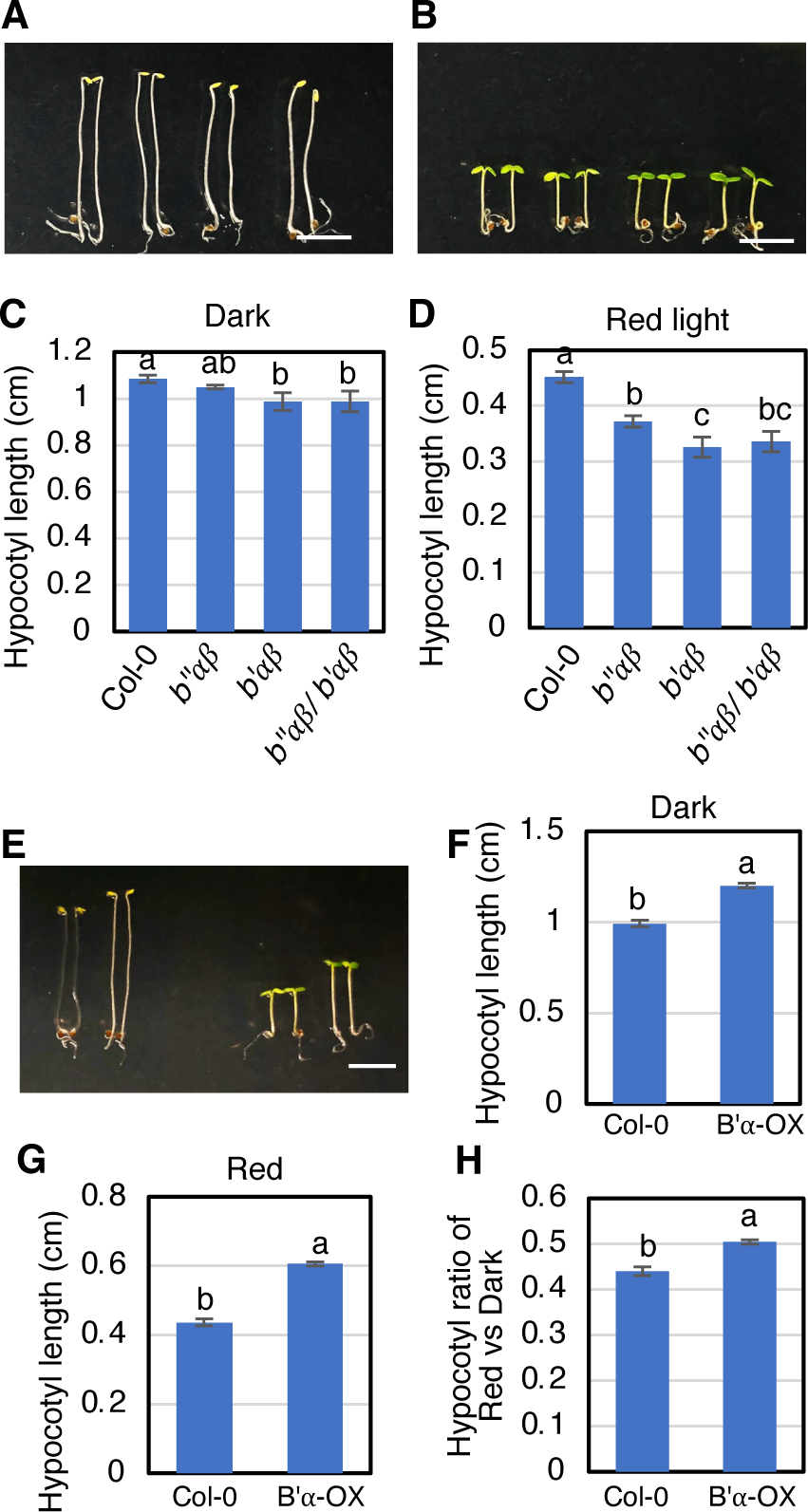
PP2A B’*α* and B’*β* promote hypocotyl elongation in red-light condition. **A and B.** Photographs showing the seedling phenotypes of *pp2ab’αβ* and *b’’αβb’αβ* grown in darkness (A) and red-light (8 μmol/m^2^s) condition (B) respectively for 4 days. The seedling order in image from left to right: Col-0, *pp2ab’’αβ*. *pp2ab’αβ* and *pp2ab’’αβb’αβ*. Scale bar in A and B: 5 mm. **C and D.** Bar graphs show the hypocotyl lengths of seedlings shown in A and B. (*n* ≥ 18). Error bars represent SE. One-way ANOVA was performed. Statistically significant differences are indicated by different lowercase letters (*P* < 0.05). **E.** Photographs showing the seedling phenotypes of B’*α* overexpression line (B’*α*-OX) grown in darkness and red-light condition (8 μmol/m^2^s) respectively for 4 days. The seedling order in image from left to right: Col-0, *B’α-OX* in dark condition (left) and red-light condition (right). Scale bar in E and F: 5 mm. **F and G.** Bar graphs show the hypocotyl lengths of seedlings shown in E. (*n* ≥30). **H.** Bar graph shows the hypocotyl length ratio of red vs dark conditions. Error bars represent SE. One-way ANOVA was performed. Statistically significant differences are indicated by different lowercase letters (*P* < 0.05).

**Supplemental Figure S8:**
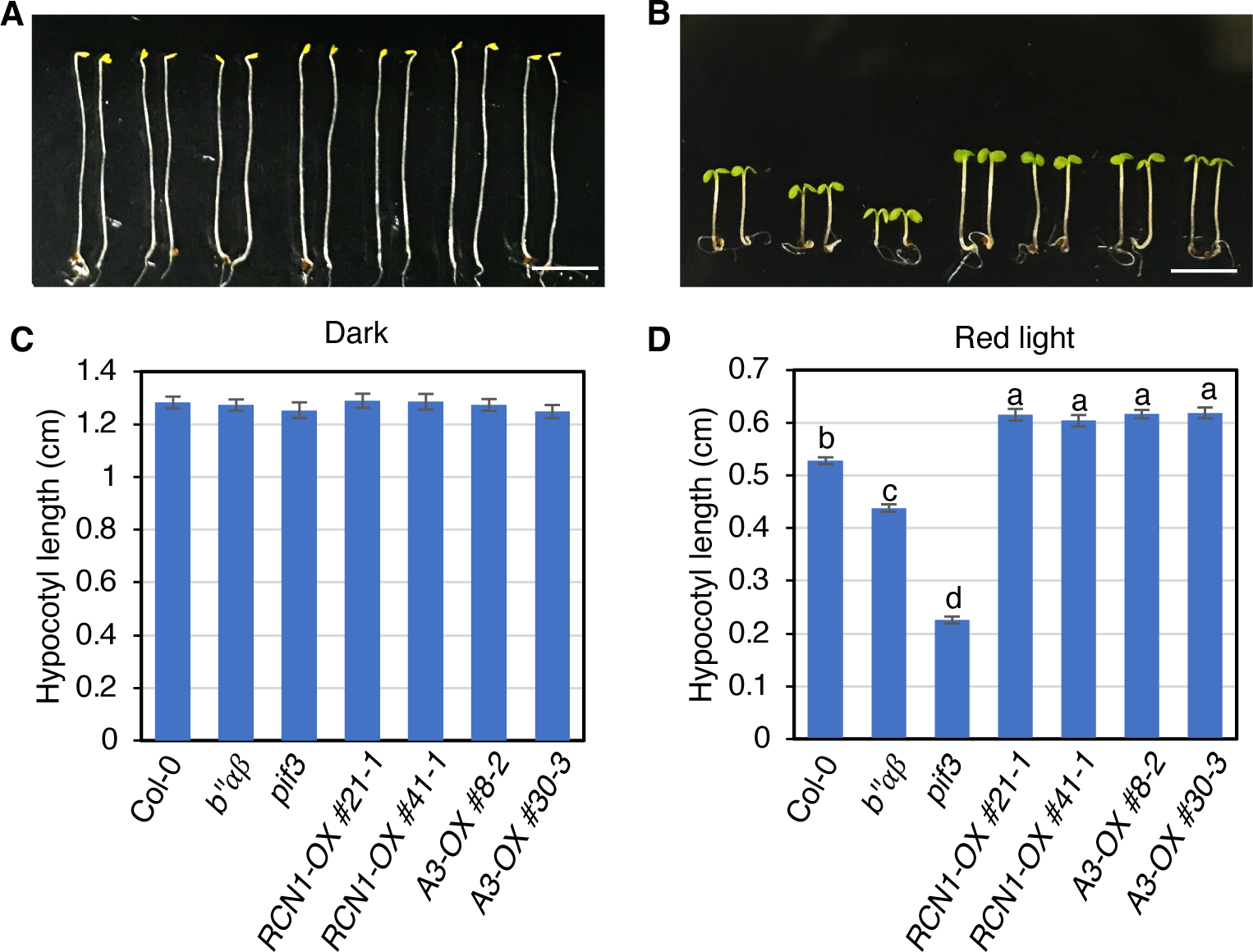
PP2A RCN1 and A3 overexpression seedlings show longer hypocotyls in red-light condition. **A and B.** Photographs showing the seedling phenotypes of *RCN1-OX/Col-0* and *A3-OX/Col-0* grown in darkness (A) and red-light (8 μmol/m^2^s) condition (B) respectively for 4 days. Col-0, *b’’αβ* and *pif3* as controls. The seedling orders in image from left to right: Col-0, *pp2ab’’αβ, pif3, RCN1-OX/Col-0 #21-1, RCN1-OX/Col-0 #41-1, A3-OX/Col-0 #8-2* and *A3-OX/Col-0 #30-3*. Scale bar in A and B: 5 mm. **C and D.** Bar graphs show the hypocotyl lengths of seedlings shown in A and B. (*n* ≥ 25). Error bars represent SE. One-way ANOVA was performed. Statistically significant differences are indicated by different lowercase letters (*P* < 0.05).

**Supplemental Figure S9:**
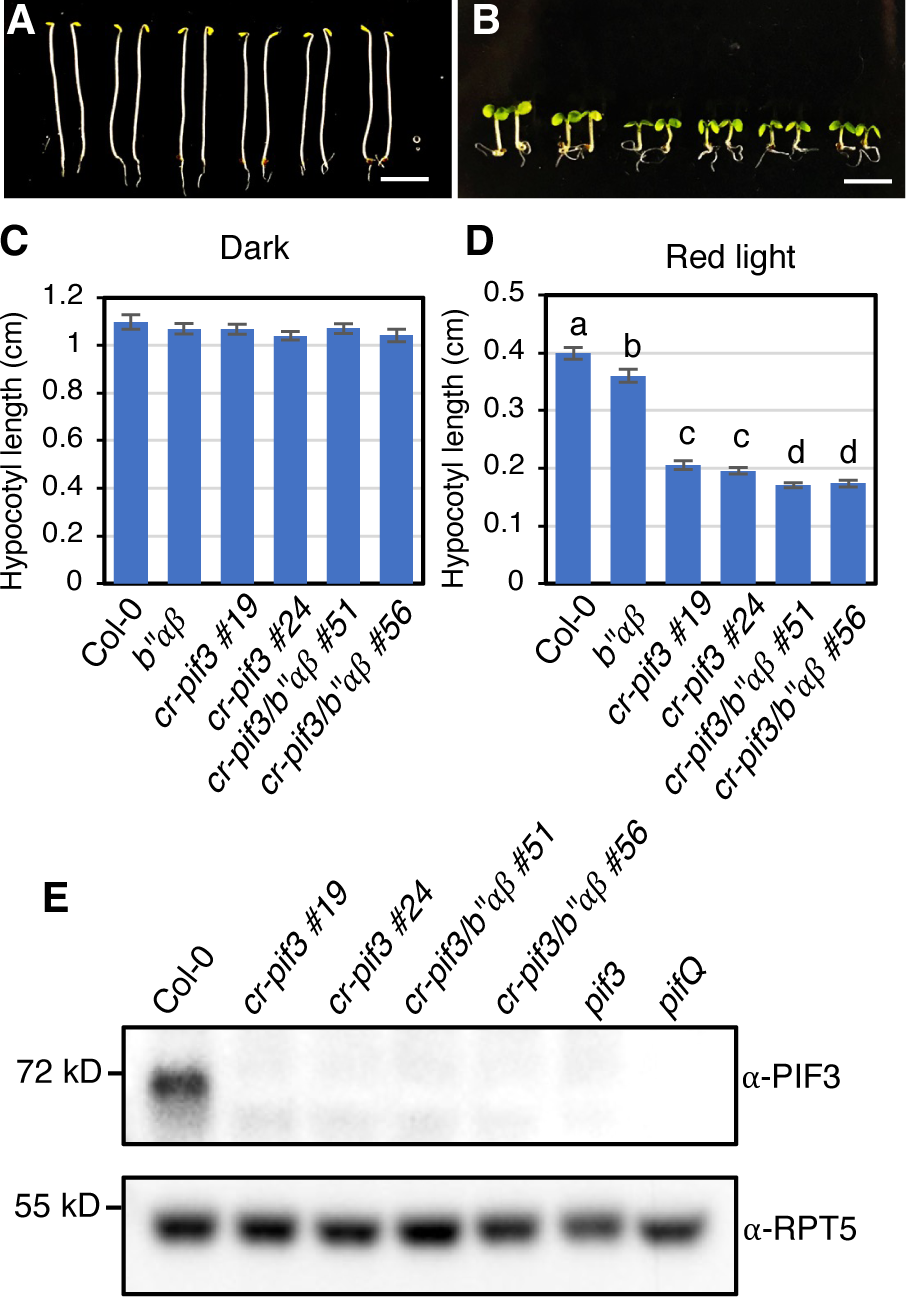
Validation of *cr-pif3* lines and phenotypic analyses under dark and red light. **A and B.** Photographs showing the seedling phenotypes grown in darkness(A) and red-light condition (B, 8 μmol/m^2^s) respectively for 4 days. The seedling order in image from left to right: Col-0, *pp2ab’’αβ, cr-pif3 #19 and #24, cr-pif3/b’’αβ #51 and #56.* Scale bar in A and B: 5 mm. **C and D.** Bar graphs show the hypocotyl lengths of seedlings shown in A and B. (*n* ≥ 19). Error bars represent SE. One-way ANOVA was performed. Statistically significant differences are indicated by different lowercase letters (*P* < 0.05). **E.** Immunoblots showing the native PIF3 protein levels in Col-0, *cr-pif3 #19 and #24, cr-pif3/b’’αβ #51 and #56. pif3* and *pifQ* as negative controls. Four-day-old dark-grown seedlings were used for protein extraction and RPT5 blot as loading control.

**Supplemental Figure S10:**
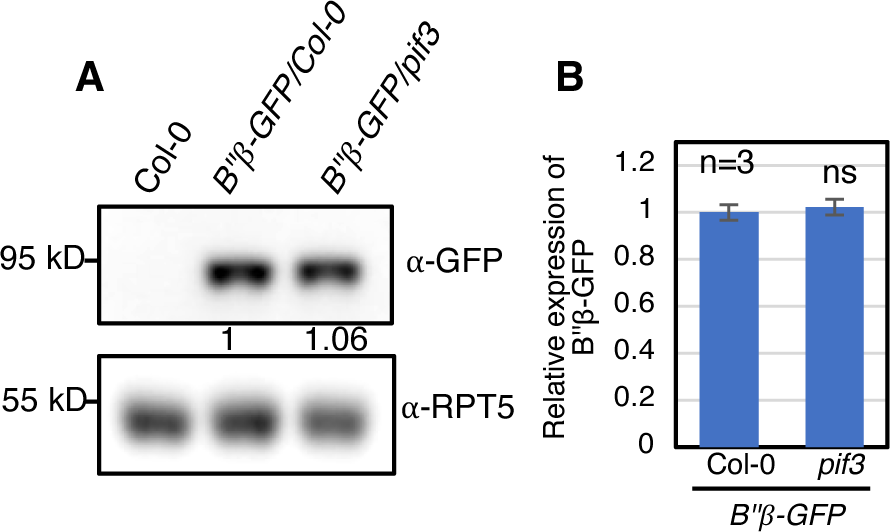
Characterization of B’’ overexpression lines. **A.** Immunoblots showing the B’’*β*-GFP protein levels in *B’’β-GFP/Col-0* and *B’’β-GFP /pif3.* Col-0 as as negative control. Four-day-old dark-grown seedlings were used for protein extraction and RPT5 blot as loading control. The numbers show the B’’*β*-GFP protein abundance after calibrating with PRT5 bands respectively. **B.** Quantification of B’’*β*-GFP protein intensity in immunoblots showed in (B). Student’s t-test was performed. Error bars represent SE (n=3).

**Supplemental Figure S11.**
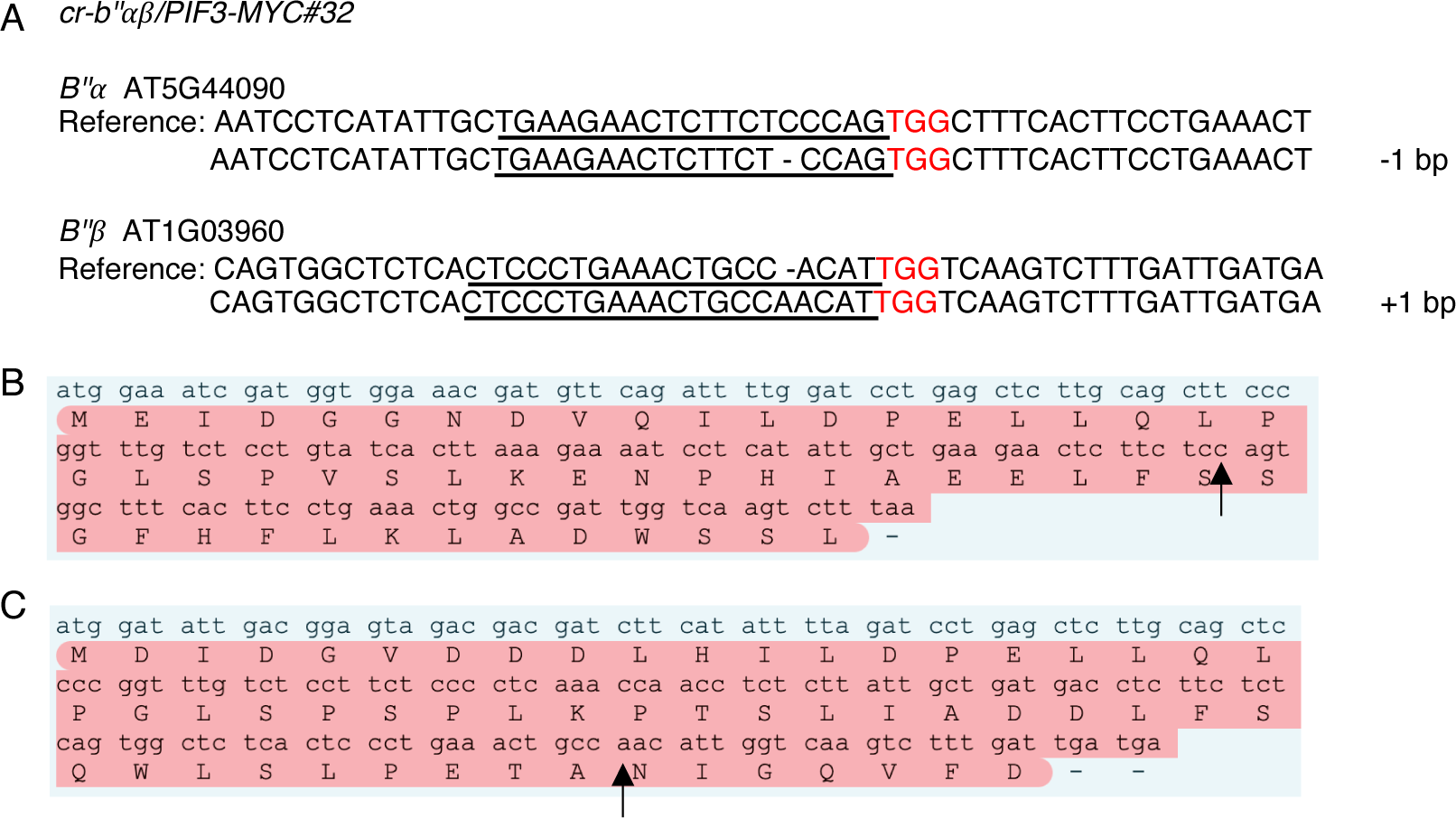
Mutation sites in *cr-b’’αβ /PIF3-MYC#32*. Mutant was generated by the CRISPR/Cas9 gene editing system (Wang and Chen, 2020). **A.** The schematic diagram of the DNA sequences of *b’’α and b’’β* CRISPR mutant in #32. Two small-guide RNAs each were designed to knock out *B’’α* and *B’’β*. The PAM (protospacer adjacent motif) sequences are marked in red. The sgRNA sequences (19 bp) are underlined. “+” sign indicates insertion and “-” sign indicates deletion. **B and C** show the translation of coding sequence *B’’α* and *B’’β* from #32 line respectively. In #32 line, the insertion in *B’’α* and deletion in *B’’β* both cause frame shift and early termination. The black arrows indicate the insertion or deletion sites.

**Supplemental Figure S12.**
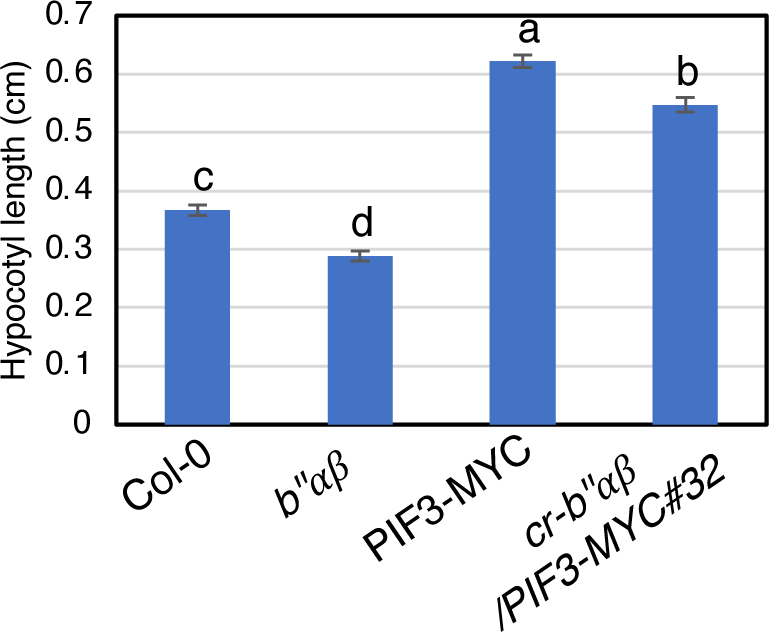
*cr-b’’αβ/PIF3-MYC#32* seedlings show short hypocotyl compared to Col-0 in red-light condition. Bar graphs show the hypocotyl lengths of seedlings of Col-0, *pp2ab’’αβ, PIF3-MYC and cr-b’’αβ/PIF3-MYC#32* grown in red-light condition (8 μmol/m^2^s) respectively for 4 days. Col-0, *pp2ab’’αβ* and PIF3-MYCas controls (*n* ≥ 25). Error bars represent SE. One-way ANOVA was performed. Statistically significant differences are indicated by different lowercase letters (*P* < 0.05).

**Supplemental Figure S13.**
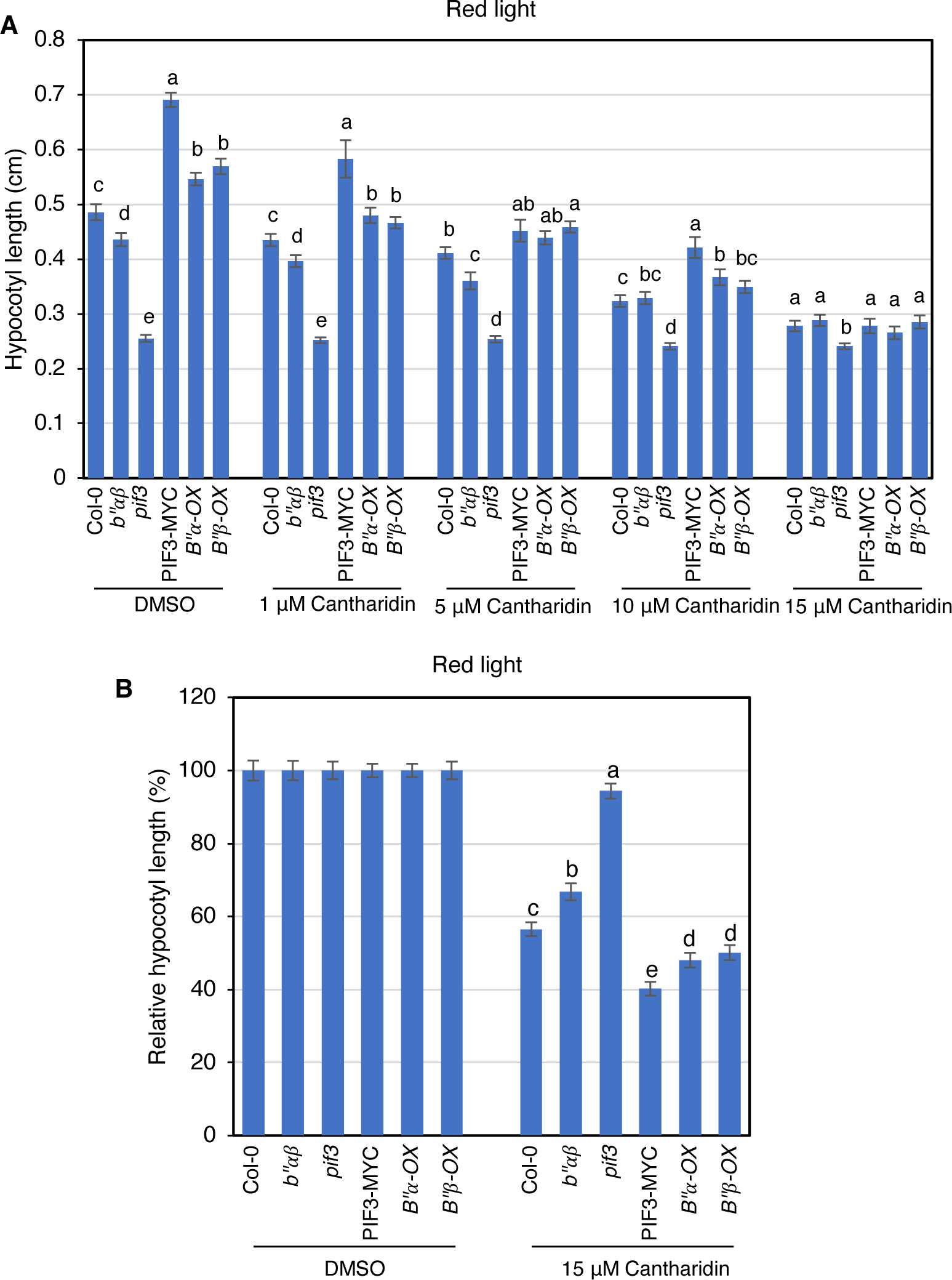
Impact of Cantharidin on Col-0, *pp2ab’’αβ, pif3*, PIF3-MYC, B’’α-OX/Col-0 and B’’β-OX/Col-0 hypocotyl elongation in red-light condition. **A.** Bar graphs show the hypocotyl lengths of seedlings of Col-0, *pp2ab’’αβ, pif3*, PIF3-MYC, *B’’α-OX/Col-0 and B’’β-OX/Col-0* grown in red-light condition (8 μmol/m^2^s) on different cantharidin as shown in the figure for 4 days. (*n* ≥ 17). **B.** Bar graph shows the relative hypocotyl lengths changes in (A) between DMSO and 15 μM cantharidin conditions. Hypocotyl lengths of seedlings from DMSO plates for different genotypes were set as 1 respectively, and then hypocotyl lengths of seedlings from 15 μM cantharidin plates were expressed relative to their DMSO controls. (*n* ≥ 17). Error bars represent SE. One-way ANOVA analysis was performed. Statistically significant differences are indicated by different lowercase letters (*P* < 0.05).

**Supplemental Figure S14:**
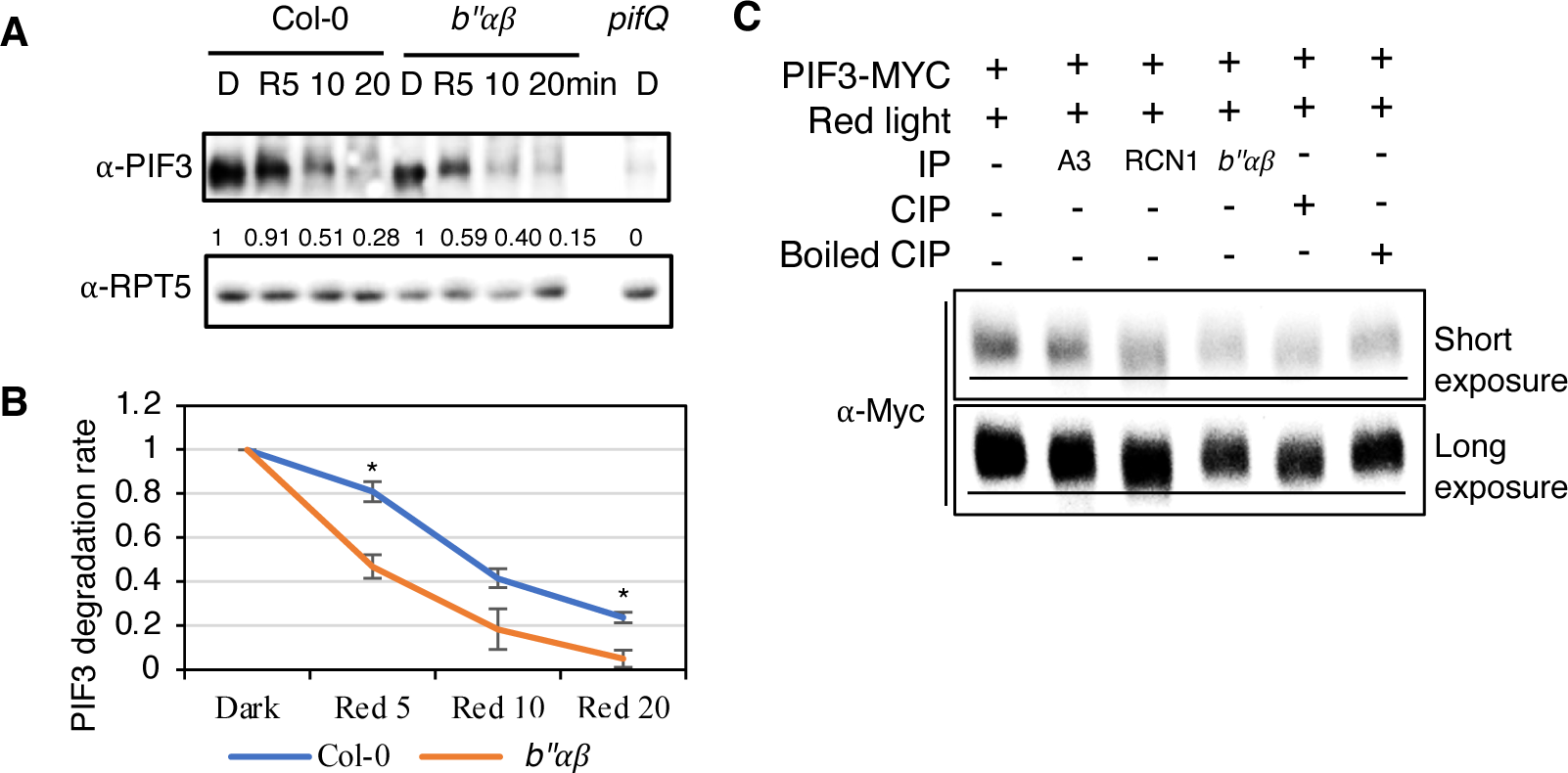
B” α and β suppress PIF3 degradation by directly dephosphorylating PIF3 under red light. **A.** Immunoblots showing the light-induced degradation of native PIF3 in the *pp2ab’’αβ* mutant compared to wild type. Four-day-old dark-grown seedlings were either kept in darkness or exposed to red light (25 μmol/m^2^s) for the duration indicated before being sampled for protein extraction. RPT5 blot as loading control. The numbers show the native PIF3 protein abundance after calibrating with PRT5 bands. **B.** Line graphs show native PIF3 degradation rate after red light exposure in wild type and *pp2ab’’αβ* mutant background based on three independent blots. * P<0.05 based on Student’s t-test. Error bars represent SE (n=3). **C.** Dephosphorylation assay was performed by using immunoprecipitated PIF3-MYC and PP2A proteins from PIF3-MYC plants and RCN1-GFP, A3-GFP transgenic plants respectively. PIF3-MYC proteins from four dark-grown seedlings of PIF3-MYC, treated with 100 μM Bortezomib for 4 hours in darkness and exposed to red-light before the immunoprecipitation. RCN1-GFP and A3-GFP immunoprecipitation (IP) products as PP2A phosphatase incubate with immunoprecipitated PIF3-MYC for 1 hour at 30℃. CIP as positive control. Boiled CIP and IP product from *pp2ab’’αβ* as negative controls. CIP and Boiled CIP treatment were performed at 37 ℃ for 1 hour. Western-blot analyses was performed with anti-Myc on SDS-PAGE.

**Supplemental Figure S15:**
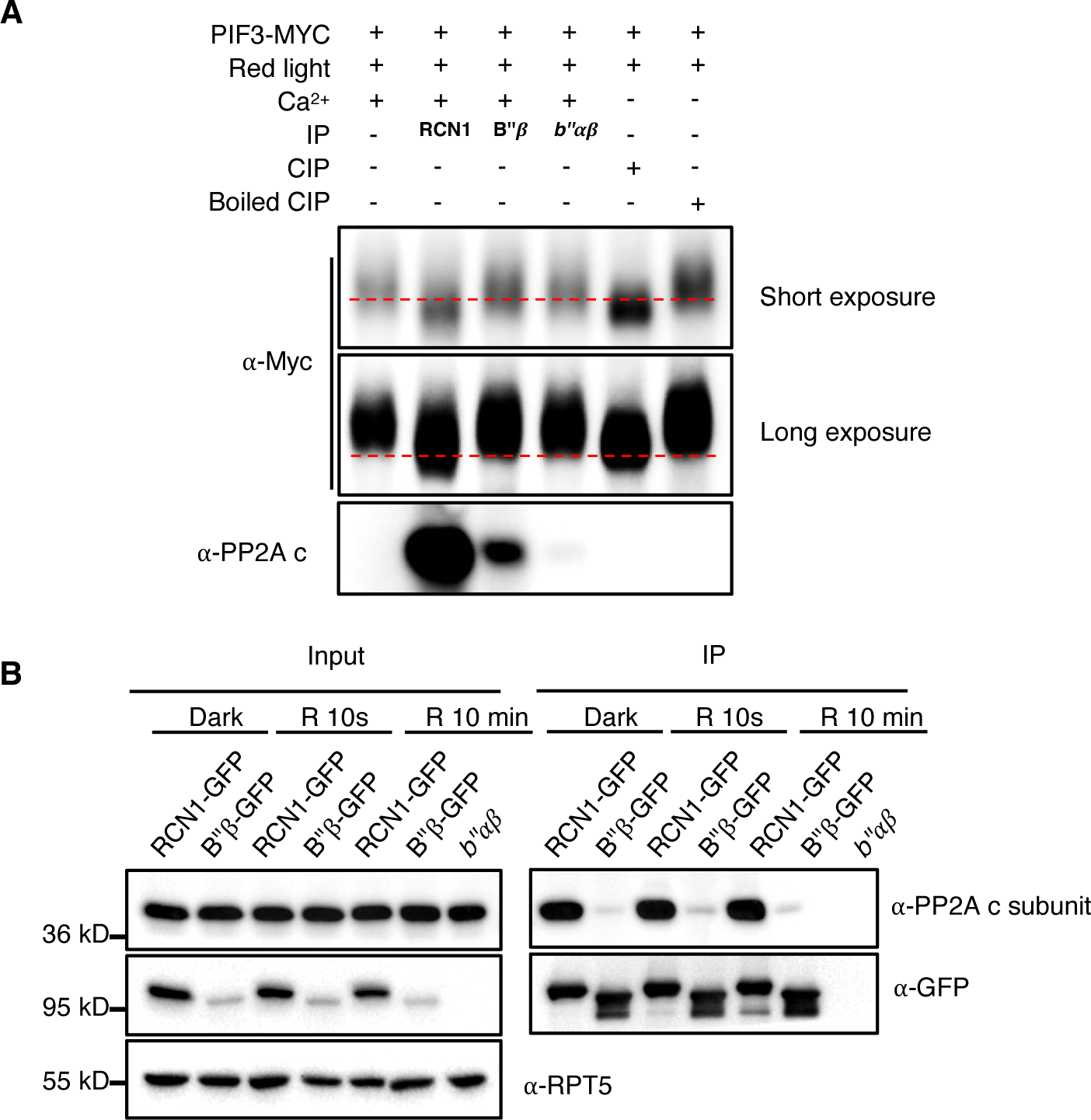
B’’*β*-GFP fails to dephosphorylate PIF3-MYC *in vitro* and exhibits lower binding ability with PP2A c subunit compared with RCN1-GFP. **A.** Dephosphorylation assay was performed by using immunoprecipitated PIF3-MYC and PP2A proteins from PIF3-MYC plants and RCN1-GFP, B’’*β*-GFP transgenic plants respectively. PIF3-MYC proteins from four dark-grown seedlings of PIF3-MYC, treated with 100 μM Bortezomib for 4 hours in darkness and exposed to red-light before the immunoprecipitation. RCN1-GFP, B’’*β*-GFP immunoprecipitation (IP) products as PP2A phosphatase incubate with immunoprecipitated PIF3-MYC for 1 hour at 30℃. Compared to RCN1-GFP IP products, double amount of B’’*β*-GFP IP products were used in this assay. CIP as positive control. Boiled CIP and IP product from *pp2ab’’αβ* as negative controls. CIP and Boiled CIP treatment were performed at 37 ℃ for 1 hour. Western-blot analyses was performed with anti-Myc on SDS-PAGE. **B.** Left panel: Immunoblots showing PP2A c subunit and GFP signals in the RCN1-GFP and B’’*β* -GFP plants. *pp2ab’’αβ* as control. Four-day-old dark-grown seedlings were either kept in darkness or exposed to red light (20 μmol/m^2^s) for the duration indicated before being sampled for protein extraction. RPT5 blot as loading control. Right panel: Immunoblots showing PP2A c subunit and GFP signals after immunoprecipitation by α-GFP and then PP2A c subunit signals were detected by α-PP2A c subunit.

**Supplemental Figure S16:**
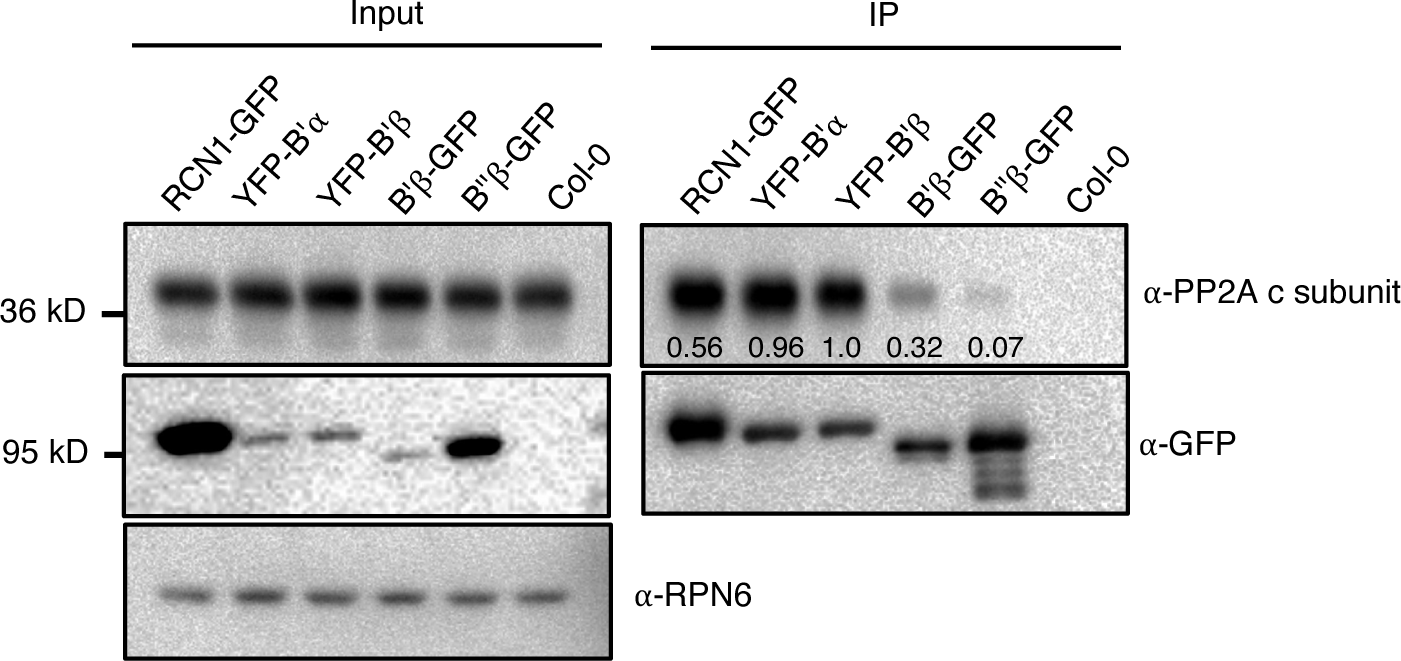
A free C-terminus is essential for B subunits to bind PP2A catalytic subunit strongly. Left panel: Immunoblots showing PP2A c subunit and GFP signals in the RCN1-GFP, YFP-B’α,YFP-B’*β*, B’*β*-GFP and B’’*β*-GFP plants. Col-0 as control. Four-day-old dark-grown seedlings were used for protein extraction. RPN6 blot as loading control. Right panel: Immunoblots showing PP2A c subunit and GFP signals after immunoprecipitation by α-GFP and then PP2A c subunit signals were detected by α-PP2A c subunit. The numbers in the right panel represent the binding ability to c subunit, generated by using c subunit signals divided by GFP signals.

**Supplemental Figure S17:**
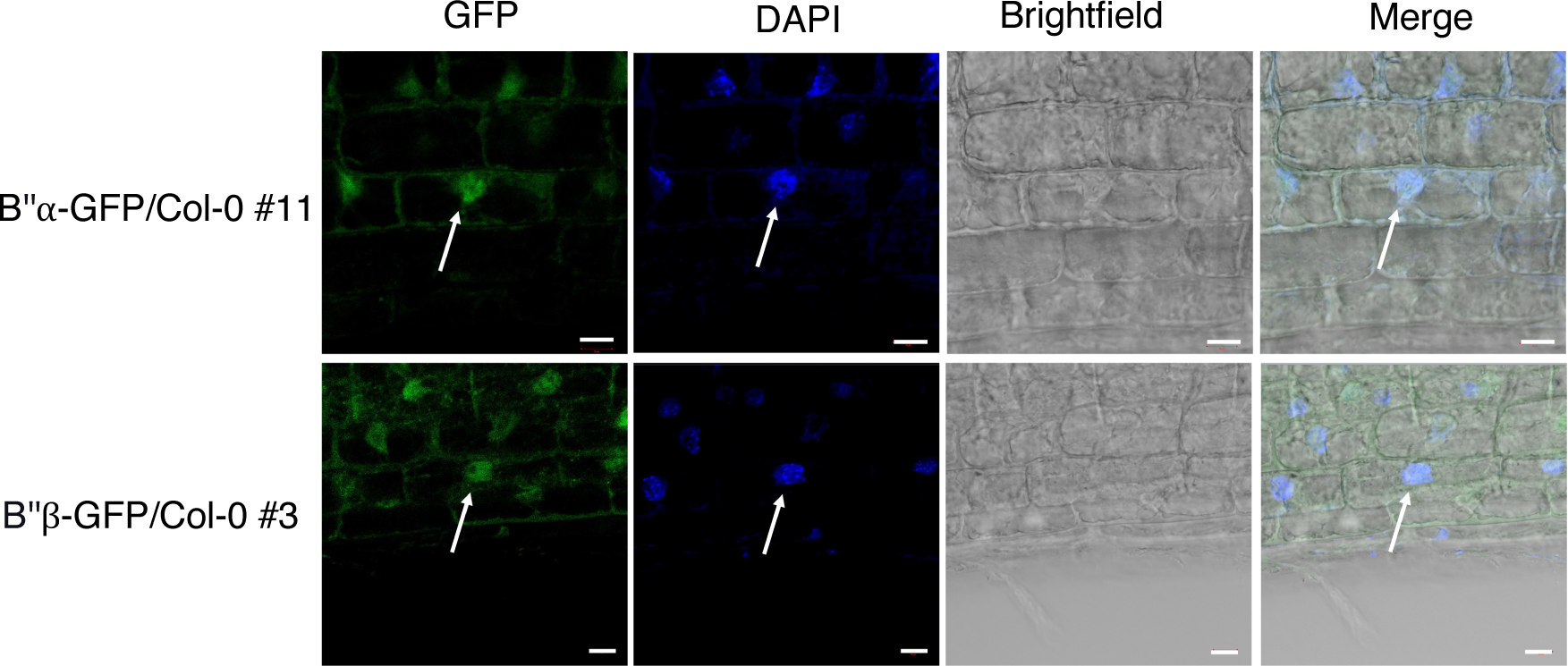
Confocal images showing the subcellular localization of B’’α-GFP (upper pane) and B’’β-GFP (lower pane) in the primary root of 4-days old seedlings grown on MS medium in the white light condition. DAPI was used to show the nucleus. White arrows show the nucleus. Scale bar is 10 μm.

**Supplemental Figure S18:**
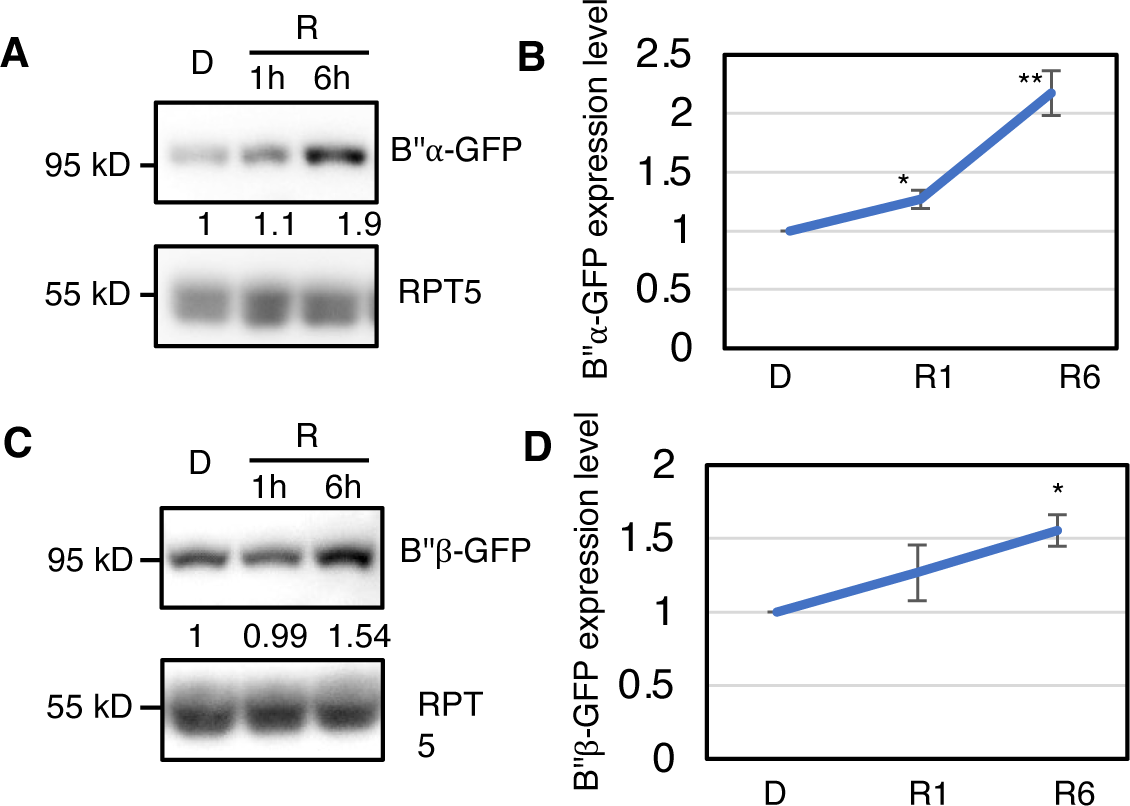
PP2AB’’*α* and B’’*β* protein levels were induced by red-light. Immunoblots showing the red-light induces expressions of B’’*α*-GFP (A) and B’’*β*-GFP (B). Four-day-old dark-grown seedlings were either kept in darkness or exposed to red light (20 μmol/m^2^s) for the 1 hour (R1) or 6 hours (R6) before being sampled for protein extraction. RPT5 blot as loading control. The numbers show the B’’*α*-GFP and B’’*β*-GFP protein abundance after calibrating with PRT5 bands respectively. Line graphs show expression levels of PP2AB’’*α*-GFP (C) and B’’*β*-GFP (D) after red light exposure based on three independent blots. * P<0.05 and ** P<0.01, based on Student’s t test. Error bars represent SE.

**Supplemental Figure S19:**
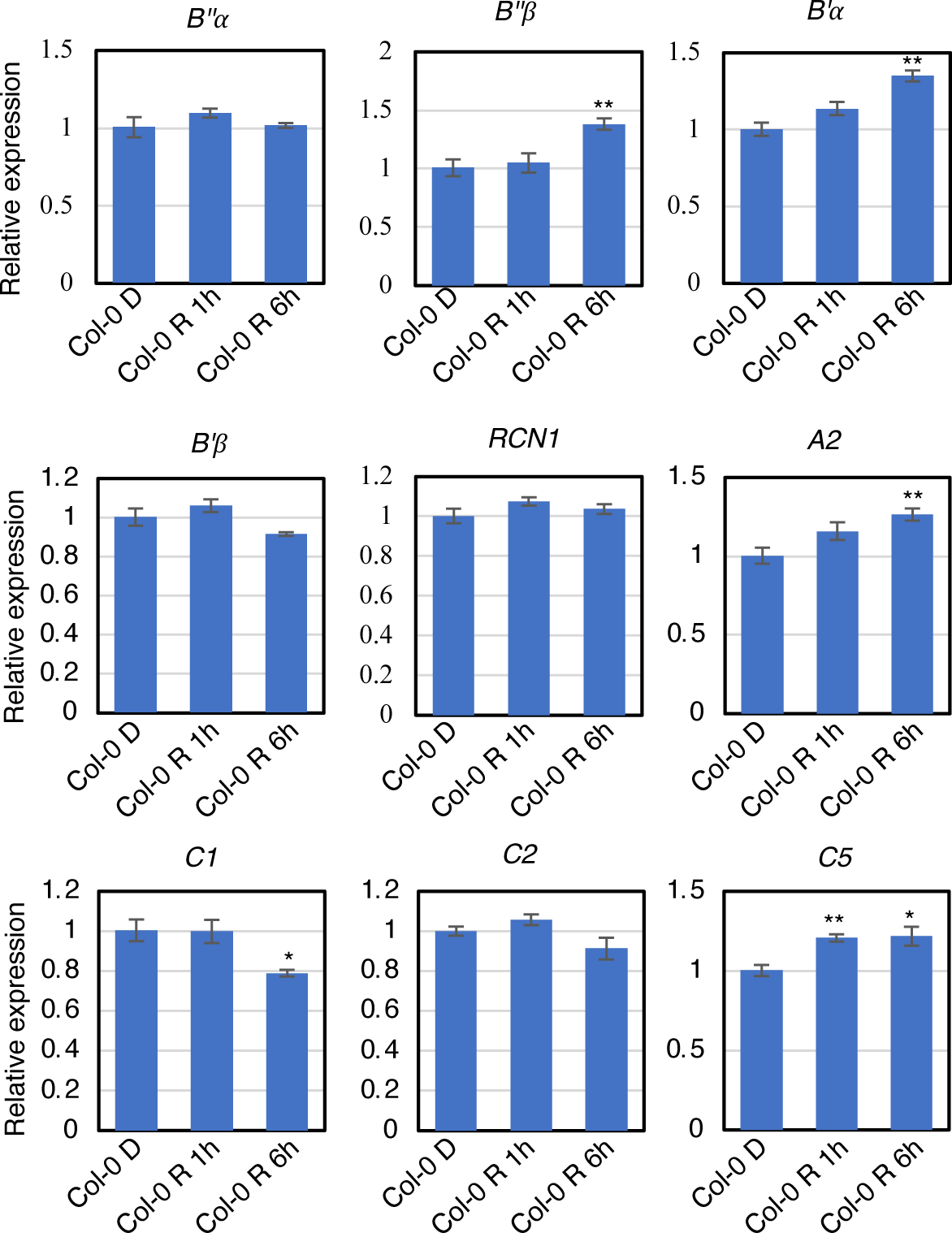
Relative expression of different subunits of *PP2A* in Col-0 after red light exposure. Total RNA was isolated from Col-0 4-day dark-grown seedlings (D) and after 1 hour (R 1h) and 6 hours (R 6h) red light (20 μmol/m^2^s) exposure. The relative expression level of *PP2A* different subunits (A subunit: *RCN1* and *A2*; B subunits: *B’’α, B’’β, B’α* and *B’β*; C subunit: *C1, C2* and *C5*) were quantified by RT-qPCR. * P<0.05 and ** P<0.01, based on Student’s t-test. Error bars represent SE ± SD (n = 4).

**Supplemental Figure S20:**
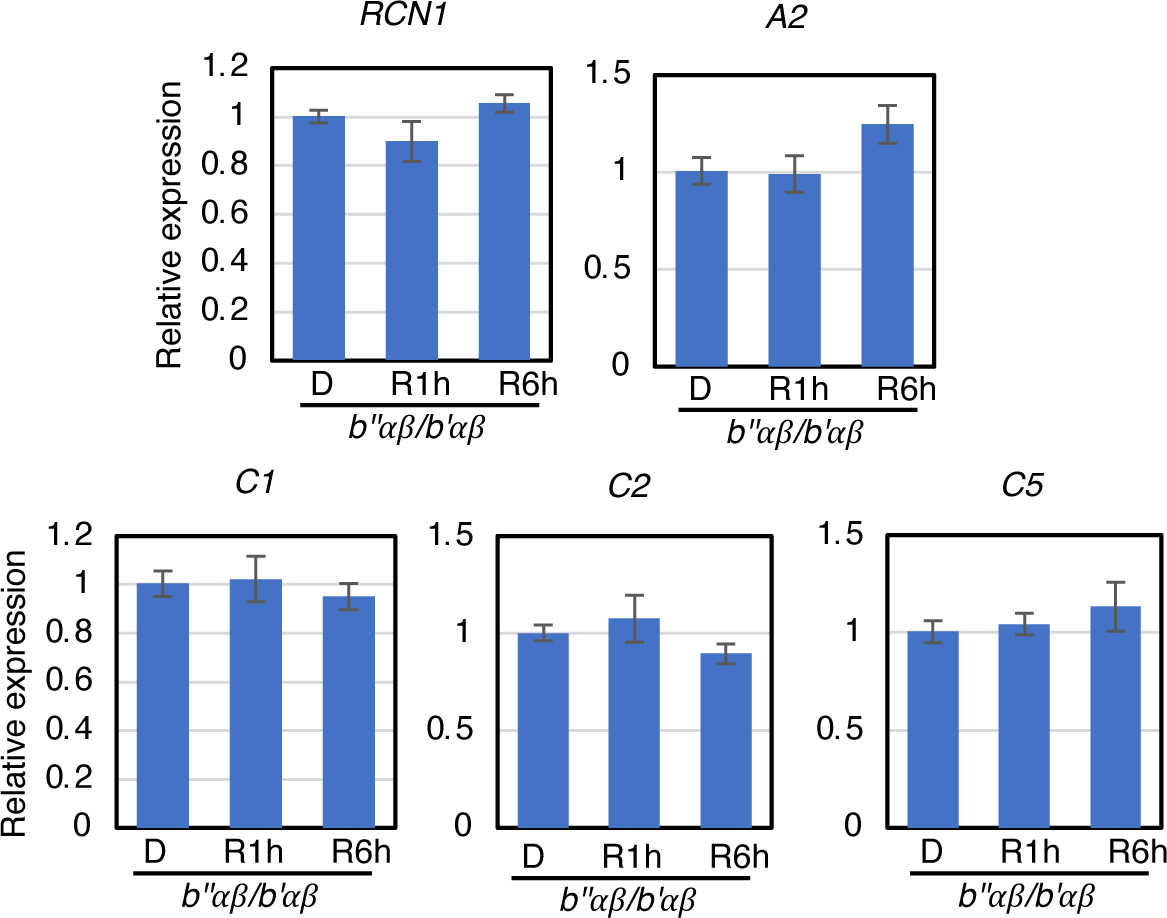
Relative expression of *PP2A A* subunits and *C* subunits in *b’’αβ/b’αβ* after red light exposure. Total RNA was isolated from *b’’αβ/ b’αβ* 4-day dark-grown seedlings (D) and after 1 hour (R1h) and 6 hours (R6h) red light (20 μmol/m^2^s) exposure. The relative expression level of PP2A different subunits (A subunit: *RCN1* and *A2*; C subunit: *C1, C2* and *C5*) were quantified by RT-qPCR. No statistical difference was found among these data based on Student’s t-test. Error bars represent SE ± SD (n = 4).

